# SCF^FBXL5^ ubiquitin ligase regulates stability of von-Hippel Lindau protein and the HIF1α-dependent response to hypoxia

**DOI:** 10.1101/2024.01.29.577840

**Authors:** Adarsh K Mayank, Vijaya Pandey, Kyle Lien, Lu Li, Yasaman Jami-Alahmadi, Ahmad Mohammad Kassem, James A Wohlschlegel

## Abstract

The canonical response to changes in cellular oxygen levels consists of the ubiquitin-dependent degradation of hypoxia-inducible transcription factors (HIFs) in a prolyl hydroxylase (PHD) and von-Hippel Lindau-ElonginB-ElonginC (VHL-ElonginBC) E3 ubiquitin ligase complex-dependent manner. This regulated degradation event is oxygen-dependent and results in activation of a transcriptional program that mediates the cellular adaptation to changes in oxygen tension. Here, we show that a distinct Cullin-RING ligase complex, SKP1-CUL1-FBXL5 (SCF^FBXL5^), physically associates with VHL and promotes its ubiquitin-dependent degradation during hypoxia. The regulation of VHL protein stability by FBXL5 influences HIF1α expression levels and the transcriptional activation of downstream hypoxia-responsive target genes. This work identifies a novel mechanism for VHL regulation which contributes to the HIF1α-mediated cellular response to hypoxia and provides an additional layer of crosstalk between iron and oxygen homeostasis.

## Introduction

Iron is a vital nutrient that serves as an essential cofactor for multiple cellular processes including aerobic respiration, protein translation, and DNA metabolism. Importantly, cells must balance the absolute requirement for iron with the oxidative stress induced toxicity associated with excess iron. The primary mechanism employed by cells to maintain intracellular iron levels within a narrow physiological window is the regulation of mRNAs encoding proteins involved in iron metabolism via a homeostatic system comprised of iron regulatory proteins 1 and 2 (IRP1/2) and the ubiquitin E3 ligase subunit FBXL5(Muckenthaler et al., 2017; Salahudeen et al., 2009; Vashisht et al., 2009). FBXL5 has the capacity to sense intracellular iron levels via iron binding to its N-terminal hemerythrin-like domain. In iron replete conditions, iron binding by FBXL5 stabilizes it, enabling it to assemble into a multi-subunit E3 ubiquitin ligase complex with SKP1, CUL1, and RBX1 (SCF^FBXL5^). SCF^FBXL5^ catalyzes the ubiquitin-dependent degradation of IRP1 and IRP2 leading to upregulation of the iron storage protein ferritin and destabilization of the transferrin receptor involved in iron uptake. During iron starvation conditions, FBXL5 is itself degraded via the ubiquitin-proteasome pathway leading to the stabilization of IRP1 and IRP2, the translational inhibition of ferritin, and the stabilization of the transferrin receptor.

In addition to its central role in iron homeostasis, FBXL5 has also been implicated in oxygen homeostasis. This is supported by several lines of evidence including (1) the regulation of IRE-containing mRNAs by IRPs is responsive to changes in oxygen tension (Rouault, 2006; Wallander et al., 2006) (2) the reversible binding of molecular O_2_ by the hemerythin-like domain of FBXL5 (Salahudeen and Bruick, 2009), and (3) the oxygen-regulated association of FBXL5 with the CIA targeting complex involved in Fe-S cluster biogenesis (Mayank et al., 2019). Although these findings broadly support the paradigm that iron and oxygen homeostatic pathways cross-regulate each other, the mechanisms by which FBXL5 can impact oxygen homeostasis are still not clearly elucidated.

In this study, we report a molecular link between FBXL5 and the key oxygen homeostatic pathway mediated by hypoxia inducible transcription factors (HIFs) and the von-Hippel Lindau (VHL)-associated E3 ubiquitin ligase complex. In this canonical pathway, HIFs are hydroxylated by the PHD family of iron-dependent prolyl hydroxylases at physiological oxygen levels, which promotes the recruitment of the VHL E3 ubiquitin ligase complex and subsequent HIF degradation (Schofield and Ratcliffe, 2004). Conversely, PHDs are inactivated in hypoxic conditions leading to HIF stabilization and the transcriptional activation of its downstream target genes. Here, we demonstrate an additional layer of regulation of the VHL-HIF axis mediated by FBXL5. Specifically, we find that SCF^FBXL5^ interacts with VHL and promotes its ubiquitin-dependent degradation. This FBXL5-dependent degradation of VHL has profound effects on both HIF1α stability and its downstream hypoxia-dependent gene expression program. Importantly, this pathway functions independently of the FBXL5-IRP axis and highlights the ability of FBXL5 to independently regulate both iron and oxygen homeostasis through the proteolysis of distinct substrates.

## Results

### FBXL5, an E3 ubiquitin ligase substrate adaptor, interacts with VHL, a substrate adaptor involved in canonical hypoxia signaling pathway

To better understand the role of FBXL5 in mediating crosstalk between iron homeostasis and other cellular processes, we identified novel FBXL5-associated proteins using an affinity purification - mass spectrometry (AP-MS) approach. Briefly, human FBXL5 was immunoprecipitated from Flp-In-293 cells stably expressing HA-FLAG-FBXL5, digested with trypsin, and analyzed by LC-MS/MS. Intriguingly, this analysis enriched not only FBXL5 and known interactors including CUL1, SKP1, and IRP2, but also the E3 ligase subunit VHL (Fig.1A). Given the prior data supporting a role for FBXL5 in coordinating iron and oxygen metabolism (Mayank et al., 2019; Salahudeen and Bruick, 2009), we sought to examine the FBXL5-VHL interaction more closely. This interaction was validated in a cell-based co-immunoprecipitation assay using HEK293 and HeLa cells transiently overexpressing 6Myc-FBXL5 and GFP-VHL (Fig. 1B, Fig. S1A). Strengthening this observation further, we found that endogenous VHL co-purified with stably expressed wild-type FBXL5 as well as FBXL5-ΔFbox, a dominant negative mutant of FBXL5 that retains its substrate binding domain but is unable to assemble with SKP1 and CUL1 (Fig.1C, Fig. S1B). Of note, the interaction of endogenous VHL with the FBXL5-ΔFbox mutant was more robust relative to wild-type FBXL5 raising the possibility of VHL being a substrate of FBXL5. The interaction between FBXL5 and VHL was additionally confirmed by both a reciprocal proteomic analysis demonstrating that FBXL5 co-purified with components of VHL-ElonginBC complex (Fig. 1D) and a cell-based co-immunoprecipitation analysis which showed the presence of YFP-FBXL5 in Myc-VHL immunoprecipitates (Fig.1E). These data establish a previously unknown interaction between FBXL5, a crucial player in cellular iron homeostasis and VHL, a conserved component of canonical cellular adaptive response to hypoxia.

**Figure 1.**
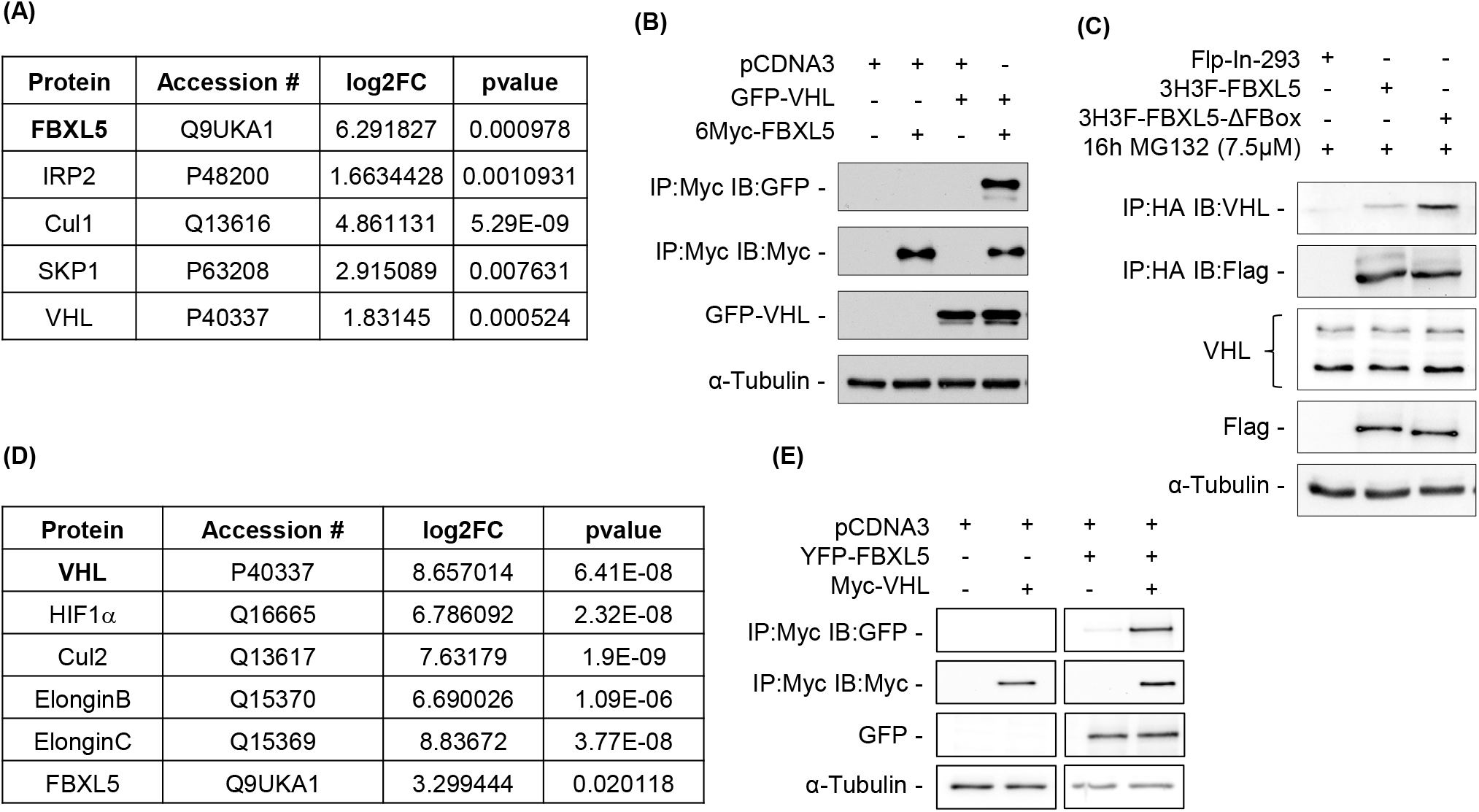
FBXL5, an E3 ubiquitin ligase substrate adaptor, interacts with VHL, a substrate adaptor involved in canonical hypoxia signaling pathway. (A) Proteins interacting with FBXL5 were identified by affinity purification coupled with mass spectrometry (AP-MS) after immunoprecipitation of HA-FBXL5 from Flp-In-293 cells stably overexpressing 3xHA-3xFLAG-FBXL5. VHL was present specifically in FBXL5 IPs but not control IPs. Log2fold-change and p-values were calculated using MSStats. (B) HEK293 cells were co-transfected with GFP-VHL and 6Myc-FBXL5. Myc-FBXL5-bound complexes were immunopurified using anti-Myc antibodies. Whole cell extracts (WCEs) and Myc-immunoprecipitates (IP:Myc) were resolved by SDS-PAGE and immunoblotted with GFP, Myc and α-tubulin antibodies. (C) Whole cell lysates from Flp-In 293 control cells or cells stably overexpressing HA-FLAG-FBXL5 or FBXL5-ΔFbox and treated with 7.5μM MG132 for 16 hours before harvesting were immunoprecipitated with anti-HA antibodies. HA immunoprecipitates (IP: HA) and WCEs were resolved by SDS-PAGE and immunoblotted with antibodies against Flag, VHL and α-tubulin. (D) Proteins interacting with VHL were identified by affinity purification coupled with mass spectrometry (AP-MS) after immunoprecipitation of HA-VHL from Flp-In-293 cells stably overexpressing 3xHA-3xFLAG-VHL. FBXL5 was specifically enriched in VHL IPs compared to control IPs. Log2fold-change and p-values were calculated using MSstats. (E) HEK293 cells were co-transfected with YFP-FBXL5 and 6Myc-VHL. Myc-VHL-bound complexes were immunopurified using Myc antibodies. Whole cell extracts (WCEs) and Myc-immunoprecipitates (IP:Myc) were resolved by SDS-PAGE and immunoblotted with antibodies against GFP, Myc, and α-tubulin.

### FBXL5 negatively regulates VHL stability by promoting its poly-ubiquitination and degradation

FBXL5 is an F-box containing substrate adaptor protein that binds to its cognate substrates IRP1 and IRP2 and brings them in proximity of SCF^FBXL5^ E3 ubiquitin ligase complex, triggering their polyubiquitination and subsequent degradation. Since FBXL5 interacts with VHL, we examined whether VHL could be a potential substrate of SCF^FBXL5^. We first tested whether the presence of FBXL5 influenced the steady state abundance of VHL. Fig. 2A shows that depletion of FBXL5 by small interfering RNAs (siRNAs) (validated using quantitative PCR, Fig. S4A) increased endogenous VHL levels relative to control siRNA consistent with the idea that FBXL5 promotes VHL degradation. Next, we measured the rate of VHL protein degradation using a cycloheximide pulse chase assay and found that the VHL was stabilized in cells depleted of FBXL5 using siRNA (Fig. 2B, Fig. S2A). Since VHL protein has been previously shown to be regulated in hypoxic cells (Turcotte et al., 2004; Wang et al., 2020), we also examined the effect of FBXL5 depletion on VHL protein stability in hypoxia. To this end, we exposed control cells and cells treated with FBXL5 siRNA to hypoxia for 16 hours, transferred them to 21% O_2_, and harvested them at indicated time points. As shown in Fig. 2C, we found that FBXL5 depletion led to stabilization of VHL protein in hypoxic cells and the effect remained intact upon exposure of cells to 21% O_2_. Together, these observations suggest FBXL5 not only interacts with VHL, but also reduces VHL protein stability in both 21 and 1% O_2_ conditions.

**Figure 2.**
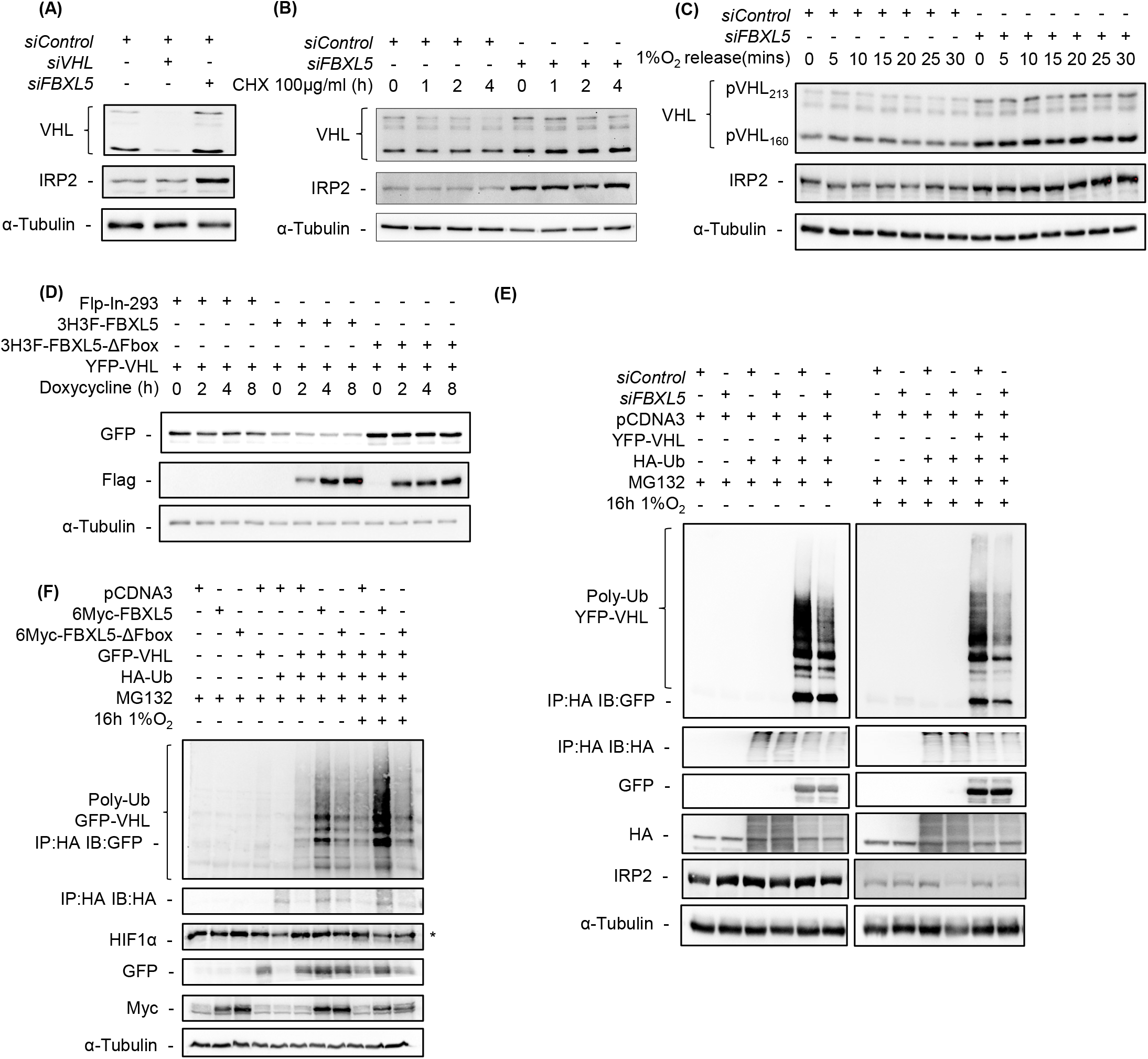
FBXL5 negatively regulates VHL stability by promoting its poly-ubiquitination and degradation. (A) HEK293 cells were treated with either non-targeting siRNA (siControl), siRNA targeted against FBXL5 (siFBXL5), siRNA targeted against VHL (siVHL) or siFBXL5 and siVHL together. WCEs were immunoblotted with VHL, IRP2 and α-tubulin antibodies. VHL siRNA was utilized in this experiment to confirm the efficacy of the VHL antibody. IRP2 stabilization was used to indicate the efficacy of siRNA-mediated depletion of FBXL5. (B) HEK293 cells were treated with either siControl or siFBXL5. Post-transfection, cells were treated with 100μg/mL of cycloheximide (CHX) to inhibit protein translation and cells were harvested at indicated time points. WCEs were immunoblotted with VHL, IRP2 and α-tubulin antibodies. IRP2 stabilization was used to indicate the efficacy of siRNA-mediated depletion of FBXL5. (C) HEK293 cells were treated with either siControl or siFBXL5. Post-transfection, cells were exposed to 1% O_2_ for 16 hours. Following re-oxygenation, cells were harvested at indicated time points. WCEs were immunoblotted with VHL, IRP2 and α-tubulin antibodies. IRP2 stabilization was used to indicate the efficacy of siRNA-mediated depletion of FBXL5. (D) Flp-In 293 cells stably expressing either HA-FLAG-FBXL5 or FBXL5-ΔFBox mutant were maintained in doxycycline-free conditions and transfected with YFP-VHL. Post-transfection, the cells were left uninduced and subjected to DFO treatment (50μM) for 16 hours to degrade FBXL5 synthesized both endogenously and as a result of leaky expression of doxycycline-inducible exogenous promoter. Following DFO treatment, cells were induced with doxycycline and harvested at indicated time points. WCEs were immunoblotted using antibodies against FLAG, GFP and α-tubulin. *denotes non-specific band in GFP panel. (E) HEK293 cells were treated with either siControl or siFBXL5 followed by co-transfection with HA-Ub and YFP-VHL. Post-transfection cells were exposed to either 21% or 1% O_2_ for 16 hours and treated with 15μM MG132 for 4 hours before harvesting. Ubiquitin conjugates were immunoprecipitated with HA antibodies. HA immunoprecipitates (IP: HA) and WCEs were immunoblotted with GFP, HA and α-tubulin antibodies. (F) HEK293 cells were co-transfected with HA-Ub and GFP-VHL along with Myc-tagged FBXL5 or FBXL5-ΔFBox mutant. Post-transfection cells were exposed to either 21% or 1% O_2_ for 16 hours and treated with 15μM MG132 for 4 hours before harvesting. Ubiquitin conjugates were immunoprecipitated by using anti-HA antibodies. HA immunoprecipitates (IP: HA) and WCEs were immunoblotted with GFP, HA, Myc, HIF1α and α-tubulin antibodies.

Given that depletion of FBXL5 stabilized VHL, we also tested if the converse was true and whether overexpression of FBXL5 would reduce VHL stability. We examined the stability of transiently overexpressed YFP-VHL in either control cells or cells stably expressing wild-type FBXL5 under the control of a doxycycline-inducible promoter. 24 hours post-transfection, we treated the cells with doxycycline to induce expression of wild-type FBXL5 and harvested them at regular time intervals to measure YFP-VHL protein levels over time as FBXL5 accumulates. As shown in Fig. 2D (lanes 5-8), there was a marked decrease in VHL levels upon expression of wild-type FBXL5 consistent with the model that FBXL5 can promote VHL degradation. We also examined the effect of overexpressing the dominant negative FBXL5-ΔFbox mutant on YFP-VHL stability (Fig 2D. Lanes 9-12) and found that VHL protein was maintained at a high level in cells overexpressing the mutant. These results indicate that FBXL5 can stimulate the downregulation of VHL protein and this activity depends upon its ability to assemble into an active SCF^FBXL5^ E3 ubiquitin ligase complex via its conserved F-box domain and is consistent with VHL as a potential substrate of SCF^FBXL5^.

In addition to examining FBXL5-dependent changes in VHL stability, we also determined if FBXL5 could influence the ubiquitination state of VHL in a manner consistent with VHL as substrate. We performed cell-based ubiquitination assays in which HEK293 cells were transfected with either control or FBXL5-specific siRNA followed by transfection with YFP-VHL and 6xHA-ubiquitin. Reduced levels of YFP-VHL in HA-immunoprecipitates confirmed that loss of FBXL5 attenuated polyubiquitination of VHL (Fig. 2E). The effect was observed both in normoxia and hypoxia treated cells. Conversely, overexpression of wild-type FBXL5 but not the FBXL5-ΔFbox mutant induced polyubiquitination of YFP-VHL in both normoxic and hypoxic cells (Fig. 2F). Since protein ubiquitination involves attachment of ubiquitin moieties to substrate lysines (Amm et al., 2014; Komander and Rape, 2012), we designed a VHL mutant, VHL-K3R, with all three lysines (K159, K171 and K196) substituted with arginines. As shown in Fig. S2B, while wild-type VHL was efficiently poly-ubiquitinated by FBXL5, VHL-K3R failed to display this effect, suggesting that the polyubiquitination of VHL induced by FBXL5 requires presence of lysines in VHL protein, in line with a classical E3 ligase-substrate relationship. These data along with the observation that endogenous VHL interacts more robustly with the FBXL5-ΔFbox substrate trapping mutant of FBXL5 (Fig. 1C) strongly argue for a role of FBXL5 in targeting VHL for ubiquitin-mediated degradation.

### VHL interacts with FBXL5 via a region spanning 114-170 amino acids

*VHL* gene encodes two mRNAs and three naturally occurring closely related protein isoforms, pVHL_213_ (longest isoform), pVHL_160_ and pVHL_172_ (Lenglet et al., 2018; Schoenfeld et al., 1998). While pVHL_172_ is a result of alternative splicing and excludes exon 2, pVHL_160_ results from translation initiation from an internal ribosome entry site. To identify the domain(s) of VHL mediating its interaction with FBXL5, we tested the ability of naturally occurring VHL isoforms as well as systematically generated deletion mutants across the VHL protein sequence (Fig. 3A) to associate with FBXL5. As shown in Fig. 3B, all the three protein isoforms of VHL interact with wild-type FBXL5 with similar robustness while a deletion mutant containing only 113 N-terminal amino acids fails to interact with FBXL5. This finding together with the observation that pVHL_160_ interacts strongly with FBXL5 suggests that the N-terminal 113 amino acids of VHL do not participate in binding with FBXL5. Interestingly, a deletion mutant of VHL comprised of exon 2 alone, encoding 114-154 amino acids, retains binding to FBXL5 indicating that the region of VHL mediating the interaction is located C-terminal to amino acid 113. We also found that the region spanning amino acids 155-189 of VHL, overlapping with ElonginBC binding box, interacted with FBXL5. However, the interaction of VHL with FBXL5 remained unaffected by deletion of the ElonginBC binding box (157-172 amino acids) in the context of full-length pVHL_213_ and pVHL_172_ isoforms (Fig. S3A), indicating that the BC box is dispensable for FBXL5 binding. Additional binding studies found that while deletion of 98-154 amino acids of VHL did not perturb its interaction with FBXL5 (Fig. S3B), two deletions mutants of VHL, Δ98-170 and Δ106-162, either lost (Δ98-170) or displayed dramatically reduced FBXL5 binding (Δ106-162) (Fig. 3C). In order to define the minimal region of VHL involved in FBXL5 binding, we generated Δ114-170 and Δ114-162 mutants and demonstrated that Δ114-162 retained binding with FBXL5 while a deletion of 8 additional amino acids (Δ114-170) was sufficient to disrupt the interaction with FBXL5 (Fig. 3D). Overall, the deletion mutagenesis studies revealed that amino acids 114-170 participate in binding of VHL to FBXL5 with the critical region required for this interaction likely housed within amino acids 163-170.

**Figure 3.**
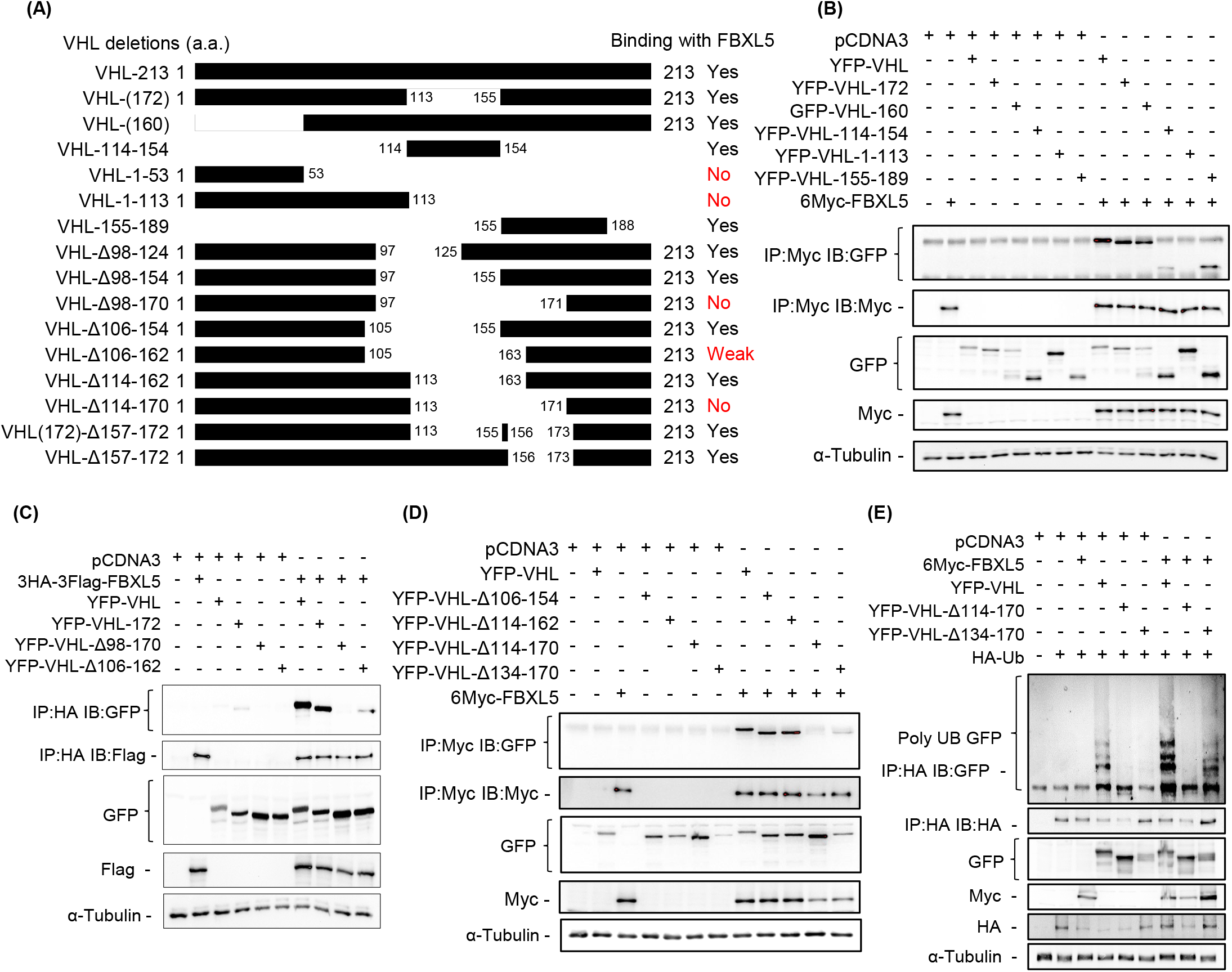
VHL interacts with FBXL5 via a region spanning 114-170 amino acids. (A) Domain structure of VHL isoforms and deletion mutants. (B) HEK293 cells were co-transfected with Myc-FBXL5 and either YFP-VHL-WT, YFP-VHL-172, GFP-VHL-160, YFP-VHL-114-154, YFP-VHL-1-113 or YFP-VHL-155-189. Whole cell extracts (WCEs) were immunoprecipitated using beads conjugated to anti-Myc antibodies. Myc-immunoprecipitates (IP:Myc) and WCEs were immunoblotted using antibodies against Myc, GFP and α-tubulin. (C) HEK293 cells were co-transfected with 3xHA-3xFlag-FBXL5 and either YFP-VHL-WT, YFP-VHL-172, YFP-VHL-Δ98-170 or YFP-VHL-Δ106-162. WCEs were immunoprecipitated using beads conjugated to anti-HA antibodies. HA-immunoprecipitates (IP:HA) and WCEs were immunoblotted using antibodies against Flag, GFP and α-tubulin. (D) HEK293 cells were co-transfected with Myc-FBXL5 and either YFP-VHL-WT, YFP-VHL-Δ106-154, YFP-VHL-Δ114-162, YFP-VHL-Δ114-170 or YFP-VHL-Δ134-170. WCEs were immunoprecipitated using beads conjugated to anti-Myc antibodies. Myc-immunoprecipitates (IP:Myc) and WCEs were immunoblotted using antibodies against Myc, GFP and α-tubulin. (E) HEK293 cells were co-transfected with HA-Ub and Myc-FBXL5 along with either YFP-VHL-WT, YFP-VHL-Δ114-170 or YFP-VHL-Δ134-170. Post-transfection, cells were treated with 15μM MG132 for 4 hours before harvesting. Ubiquitin conjugates were immunoprecipitated by using anti-HA antibodies. HA immunoprecipitates (IP: HA) and WCEs were immunoblotted with GFP, HA, Myc and α-tubulin antibodies.

The generation of VHL mutants that fail to interact with FBXL5 enabled us to determine whether the association of VHL with FBXL5 was strictly required for FBXL5-dependent ubiquitination as expected in a direct relationship. To test this idea, we overexpressed Myc-FBXL5 with YFP-VHL mutants Δ114-170 and Δ98-170 that are unable to associate with FBXL5 in HEK293 cells and used a cell-based ubiquitination assay to assess whether FBXL5 could stimulate ubiquitination of the mutants. As shown in Fig. 3E and Fig. S3C, while poly-ubiquitination of wild-type VHL was stimulated by Myc-FBXL5 overexpression, the Δ114-170, Δ98-170 and Δ106-162 mutants failed to display increased FBXL5-dependent polyubiquitination. On the contrary, Δ134-170 which retains binding to FBXL5 (Fig. 3D), was efficiently polyubiquitinated by Myc-FBXL5 overexpression (Fig. 3E, lanes 6 and 9). These observations indicate that the amino acids 114-170 are crucial for mediating the interaction of VHL with FBXL5 and their deletion not only abolishes binding to FBXL5 but also impairs FBXL5-mediated poly-ubiquitination of VHL.

### FBXL5-VHL interaction regulates HIF1α protein stability and HIF1α-mediated transcriptional activation under hypoxia

VHL protein regulates oxygen homeostasis by catalyzing the ubiquitination of the HIFα subunit of HIF transcription factors to facilitate their poly-ubiquitination and subsequent degradation (Ivan et al., 2001; Jaakkola et al., 2001; Maxwell et al., 1999). To assess the functional relevance of the FBXL5-VHL interaction, we next examined whether the FBXL5-dependent degradation of VHL impacted downstream HIF1α regulation. We monitored changes in HIF1α protein levels in response to FBXL5 down-regulation. As expected, cells exhibited induction of HIF1α protein levels in untreated cells upon exposure to hypoxia (Fig. 4A). However, sustained depletion of endogenous FBXL5 achieved using a CRISPRi transcriptional repression approach showed a marked dampening of HIF1α induction, a phenotype robustly rescued by overexpression of wild-type FBXL5. Acute suppression of FBXL5 expression in cells using RNA interference, also showed that FBXL5 depletion severely impairs HIF1α accumulation in response to hypoxia (Fig. 4B). Notably, the silencing of VHL in FBXL5-depleted cells rescued the HIF1α protein levels under hypoxia. The effectiveness of FBXL5 depletion was assessed by monitoring *FBXL5* mRNA levels (Fig. S4A). As shown in Fig. S4B, the effect of FBXL5 deficiency on HIF1α was not mediated by a transcriptional mechanism. As expected, VHL down-regulation leads to higher accumulation of HIF1α in both normoxia and hypoxia conditions compared to controls. It should be noted that the positive influence of combinatorial depletion of VHL and FBXL5 on HIF1α levels in hypoxic cells phenocopies the impact of restoration of FBXL5 expression in the FBXL5-depleted CRISPRi line.

**Figure 4.**
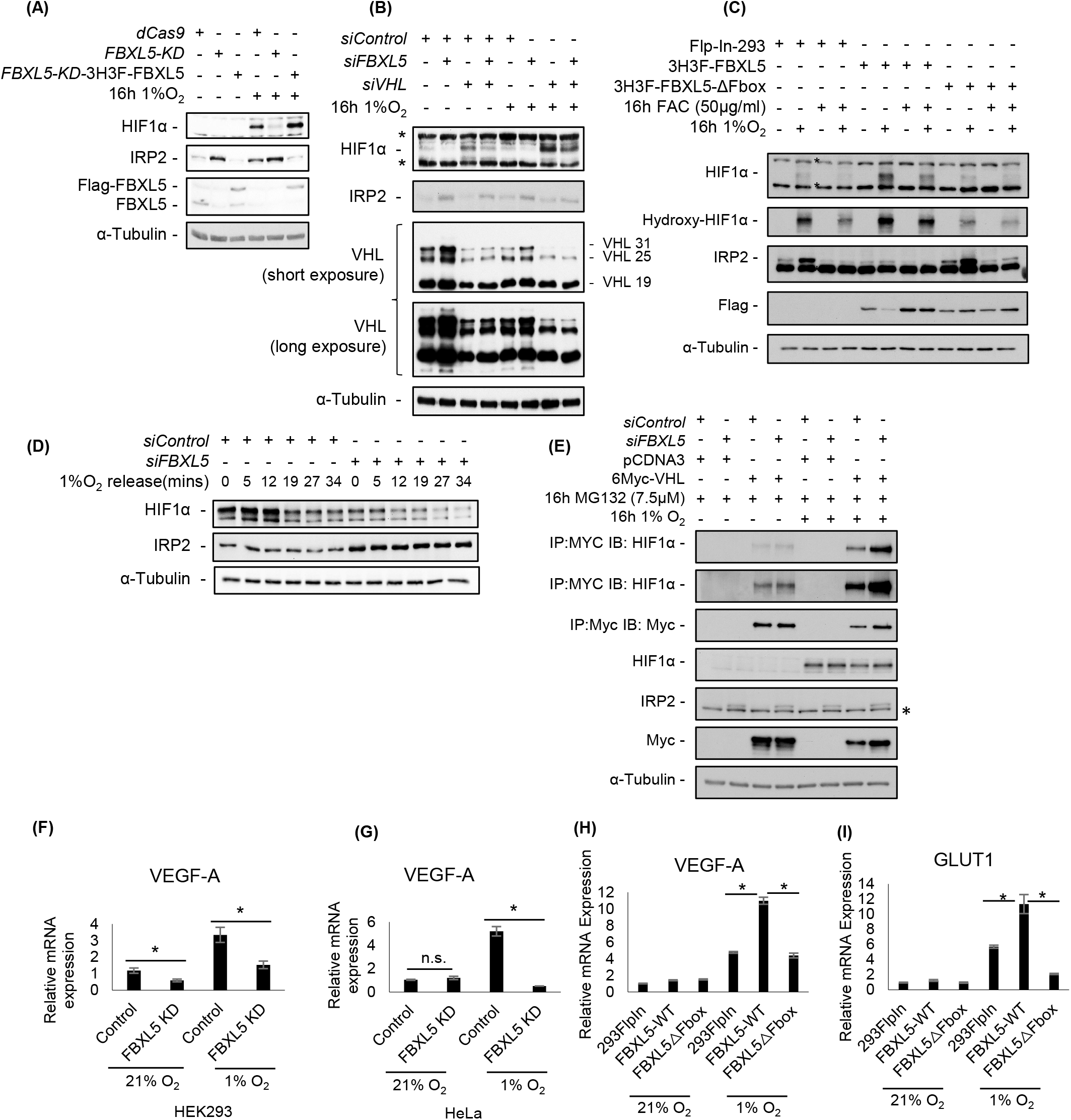
FBXL5-VHL interaction regulates HIF1α protein stability and HIF1α-mediated transcriptional activation under hypoxia. (A) Flp-In 293 cells were infected with lentiviral vectors expressing dCas9 with non-target (dCas9) or FBXL5 specific gRNA (*FBXL5* KD). *FBXL5*-KD cells lines were further manipulated to stably express HA-FLAG-FBXL5-WT (*FBXL5-KD-3H3F-FBXL5*). *dCas9, FBXL5*-*KD* and *FBXL5-KD-3H3F-FBXL5* cells were exposed to either 21 or 1% O_2_ for 16 hours. Whole cell extracts (WCEs) were immunoblotted with HIF1α, FBXL5, IRP2 and α-tubulin antibodies. (B) HEK293 cells were treated with either non-targeting siRNA (siControl), siRNA targeted against FBXL5 (siFBXL5), siRNA targeted against VHL (siVHL) or siFBXL5 and siVHL together. Post-transfection, cells were exposed to either 21 or 1% O_2_ for 16 hours. WCEs were immunoblotted with HIF1α, VHL, IRP2 and α-tubulin antibodies. IRP2 stabilization was used to indicate the efficacy of siRNA-mediated depletion of FBXL5. (C) Flp-In 293 cells stably overexpressing either HA-Flag-FBXL5 or FBXL5-ΔFBox and control cells (Flp-In 293) were treated with 50μg/mL ferric ammonium citrate (FAC) and exposed to either 21 or 1% O_2_ for 16 hours. WCEs were immunoblotted using antibodies for HIF1α, hydroxy-HIF1α, FLAG, IRP2, and α-tubulin. *denotes non-specific band in IRP2 and HIF1α panels. (D) HEK293 cells were treated with either siControl or siFBXL5. Post-transfection, cells were exposed to 1% O_2_ for 16 hours. Following re-oxygenation, cells were harvested at indicated time points. WCEs were immunoblotted with HIF1α, IRP2 and α-tubulin antibodies. IRP2 stabilization was used to indicate the efficacy of siRNA-mediated depletion of FBXL5. (E) HEK293 cells were treated with siControl or siFBXL5 followed by transfection with Myc-VHL. Post-transfection, cells were treated with 7.5μM MG132 and exposed to either 21 or 1 % O_2_ for 16 hours. Whole cell lysates were immunoprecipitated using beads conjugated to anti-Myc antibodies. Myc-immunoprecipitates (IP:Myc) and WCEs were immunoblotted using antibodies against Myc, HIF1α, IRP2 and α-tubulin. IRP2 stabilization was used to indicate the efficacy of siRNA-mediated depletion of FBXL5. *denotes non-specific band in IRP2 panel. (F-G) HEK293 (F) or HeLa-Flp-In (G) cells were treated with either siControl (Control) or siFBXL5 (FBXL5 KD). Post-transfection, cells were exposed to either 21 or 1% O_2_ for 16 hours. Relative mRNA levels of *VEGF-A* were determined using qPCR. *denotes p-value<0.05 or 0.02. (H-I) Flp-In 293 cells stably overexpressing either HA-Flag-FBXL5 or FBXL5-ΔFBox and control cells (Flp-In 293) were exposed to either 21 or 1% O_2_ for 16 hours. Relative mRNA levels of *VEGF-A* (H) and *GLUT1* (I) were determined using qPCR. *denotes p-value<0.05 or 0.02.

To further characterize the impact of FBXL5 perturbation on HIF1α protein levels, we subjected cells stably overexpressing either wild-type FBXL5 or FBXL5-ΔFbox to hypoxia. Hypoxia treatment substantially increased HIF1α levels in cells overexpressing wild-type FBXL5, but not the FBXL5-ΔFbox mutant, which is defective in promoting VHL protein degradation (Fig. 4C). Consistent with these findings, treating cells stably overexpressing wild-type FBXL5 with hypoxia mimetics such as cobalt chloride (CoCl_2_), dimethyloxalylglycine (DMOG) or the iron chelator desferroxamine (DFO) similarly led to dramatic increases in HIF1α protein levels (Fig. S4C). On the contrary, overexpression of FBXL5-ΔFbox failed to promote HIF1α protein accumulation under these conditions. Together with Fig. 4A and 4B, these observations suggest that FBXL5 positively regulates HIF1α levels in hypoxia and this function is mediated by its ability to ubiquitinate VHL.

We next determined the effect of FBXL5 depletion on HIF1α protein stability during hypoxia. Cells were treated with either control or FBXL5-specific siRNA for 24 hours, exposed to hypoxia for 16 hours, re-oxygenated, and then harvested at the indicated time points. Upon re-oxygenation, FBXL5 depleted cells displayed a faster rate of HIF1α protein destabilization compared to control cells (Fig. 4D). These observations are consistent with FBXL5 depletion stabilizing VHL and enabling its assembly into active VHL-ElonginBC complexes, thus promoting HIF1α degradation. In agreement with this idea, FBXL5-deficient cells exhibited increased accumulation of VHL protein upon re-oxygenation relative to controls, suggesting that FBXL5 depletion positively impacts VHL protein stability and VHL-dependent degradation of HIF1α (Fig. 2C). We next tested the ability of VHL to bind HIF1α in FBXL5-depleted cells exposed to hypoxia. As shown in Fig. 4E, Myc-VHL binds more robustly to HIF1α in FBXL5-depleted cells compared to control cells. Importantly, the enhanced ability of VHL to recruit HIF1α protein for degradation in FBXL5-deficient cells could not be attributed to changes in the ability of VHL to assemble with other components of the VHL-ElonginBC complex as we find that Cul2 co-purifies with VHL immunoprecipitates robustly in both control and FBXL5-deficient cells (Fig. S4D). Together these observations suggest that FBXL5 depletion leads to stabilization of VHL and increased degradation of HIF1α.

HIF1αβ heterodimer functions as a transcription factor that binds to cognate HREs in the mRNAs of hypoxia-responsive genes to regulate their expression (Semenza, 1996, 2009). To determine how FBXL5 depletion impacts the hypoxia transcriptional program, we assessed changes in HIF-dependent transcriptional activation by measuring changes in the mRNA levels of canonical HIF target genes, *VEGF-A* and *GLUT1*, during FBXL5 deficiency. As shown in Fig. 4F and 4G, HEK293 and HeLa cells depleted of FBXL5 display reduced HIF1-mediated transcriptional activation of *VEGF-A*. Conversely, overexpression of wild-type FBXL5 leads to robust induction of *VEGF-A* and *GLUT-1* genes compared to control cells undergoing hypoxia, an effect which overexpression of the FBXL5-ΔFbox mutant fails to elicit (Fig. 4H and 4I). Therefore, FBXL5 is a positive regulator of HIF1α transcriptional activity which can likely be attributed to its ability to interact with and target VHL to SCF^FBXL5^ mediated ubiquitination and degradation.

### SCF^FBXL5^ complex employs distinct mechanisms for regulation of VHL and IRP2 protein stability

Recent work has shown that FBXL5 binds to a [2Fe2S] cluster using four strictly conserved cysteine residues at its C-terminus (Cys 662, 676, 686 and 687, substituted to serine in FBXL5-4C>S mutant) and that this binding strongly influences its interaction with IRP2 (Wang et al., 2020). We next examined whether Fe-S cluster binding by FBXL5 similarly influences VHL binding. We tested the previously reported FBXL5-4C>S mutant and FBXL5-Δ666-691 mutant designed in the present study (Fig. S5A), lacking three of the conserved cysteine residues along with the previously reported interface and lid loops critical for cofactor binding, both of which fail to integrate [2Fe2S] cluster and lose interaction with IRP2 (Fig. 5A and Fig. S5B), for their ability to bind to VHL. We find that VHL interacts with wild-type FBXL5 as well as the FBXL5-4C>S and FBXL5-Δ666-691 mutants (Fig. 5A and Fig. S5D), unlike IRP2 which fails to interact with both the mutants (Fig. 5A and Fig. S5B), a phenotype reiterated in the proteomics analysis of wild-type FBXL5 and 4C>S mutant (Fig. 5B). These observations prompted us to examine and compare the poly-ubiquitination levels of VHL and IRP2 in HEK293 cell lines overexpressing either wild-type FBXL5, the FBXL5-4C>S or FBXL5-Δ666-691 mutant. As shown in Fig. 5D and Fig. S5C, FBXL5-4C>S and FBXL5-Δ666-691 mutants fail to poly-ubiquitinate Flag-IRP2 relative to wildtype FBXL5 consistent with their inability to bind IRP2. In contrast, both the [2Fe2S] cluster deficient mutants retain the ability to promote poly-ubiquitination of YFP-VHL, albeit to a lesser extent compared to wild-type FBXL5 (Fig. 5E and Fig. S5E). Consistent with this observation, HIF1α protein levels are high under hypoxia in cells stably expressing both wild-type as well as 4C>S mutant, relative to controls, indicating comparable ability of wild-type and 4C>S mutant to promote VHL degradation (Fig. 5C). Based on these findings, we conclude that FBXL5 binding to a [2Fe2S] cofactor is dispensable for VHL binding and degradation, suggesting that SCF^FBXL5^ complex employs different mechanisms for mediating IRP and VHL degradation and that the FBXL5-VHL axis can function independently of the FBXL5-IRP pathway.

**Figure 5.**
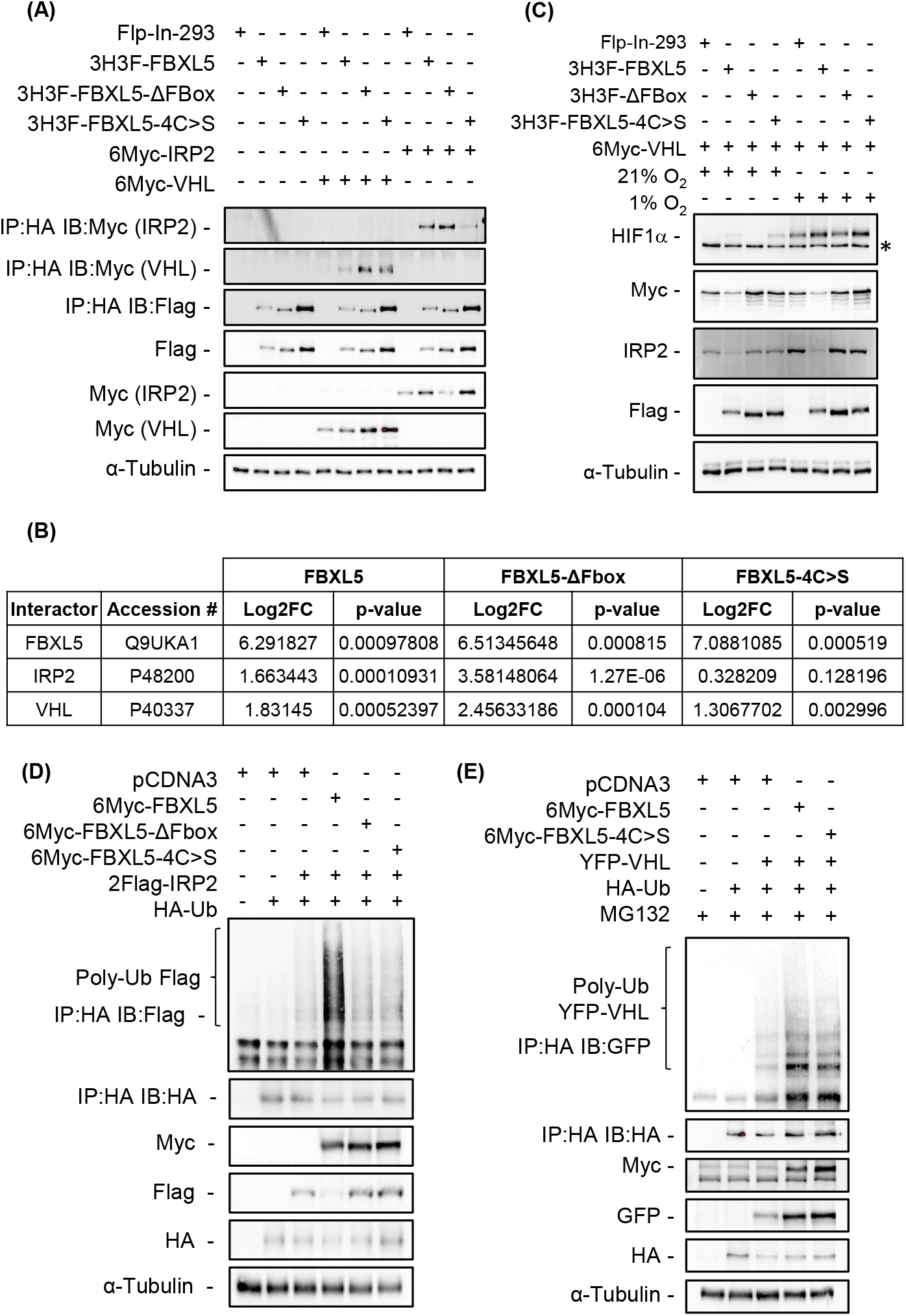
SCF^FBXL5^ complex employs distinct mechanisms for regulation of VHL and IRP2 protein stability. (A) Flp-In 293 cells stably expressing either HA-FLAG-FBXL5, FBXL5ΔFBox or FBXL5-4C>S mutants and control cells (Flp-In-293) were transfected with either 6Myc-IRP2 or 6Myc-VHL. Whole cell lysates were used for immunoprecipitation using anti-HA beads. HA immunoprecipitates and WCEs were immunoblotted using antibodies specific for Flag, Myc and α-tubulin. (B) Proteins interacting with FBXL5 were identified by affinity purification coupled with mass spectrometry (AP-MS) after immunoprecipitation of HA-tagged FBXL5 from Flp-In-293 cells stably overexpressing 3XHA-3XFLAG-tagged FBXL5, FBXL5ΔFBox or FBXL5-4C>S mutant. VHL and IRP2 were found to be specifically enriched in FBXL5 and FBXL5ΔFBox IPs compared to control IPs. Additionally, VHL, but not IRP2, was found to be enriched in FBXL5-4C>S IP compared to control. Log2fold-change and p-values were calculated using MSstats. (C) Flp-In 293 cells stably expressing either HA-FLAG-FBXL5, FBXL5ΔFBox or FBXL5-4C>S mutants and control cells (Flp-In-293) were transfected with 6Myc-VHL and subjected to either 21% or 1% O_2_ for 16 hours. WCEs were immunoblotted using antibodies specific for HIF1α, Flag, Myc, IRP2 and α-tubulin. *denotes non-specific band in HIF1α panel. (D) HEK293 cells were co-transfected with HA-Ub and Flag-IRP2 along with either Myc-FBXL5-WT, FBXL5ΔFBox or FBXL5-4C>S mutant. Post-transfection cells were treated with 15μM MG132 for 4 hours before harvesting. Ubiquitin conjugates were immunoprecipitated by using anti-HA antibodies. HA immunoprecipitates (IP: HA) and WCEs were immunoblotted with Flag, HA, Myc and α-tubulin antibodies. (E) HEK293 cells were co-transfected with HA-Ub and YFP-VHL along with either Myc-FBXL5-WT or FBXL5-4C>S mutant. Post-transfection cells were treated with 15μM MG132 for 4 hours before harvesting. Ubiquitin conjugates were immunoprecipitated by using anti-HA antibodies. HA immunoprecipitates (IP: HA) and WCEs were immunoblotted with GFP, HA, Myc and α-tubulin antibodies.

## Discussion

Oxygen and iron are coordinately regulated in response to a variety of environmental signals and across a wide range of biological processes. Understanding the precise molecular mechanisms underlying the interplay between iron and oxygen homeostasis is imperative to unveil the significance of this crosstalk in health and disease. In the present study, we describe a novel physical and functional interaction between FBXL5 and VHL, two E3 ubiquitin ligase subunits that serve as master regulators for iron and oxygen homeostasis, respectively (Fig. 6). The association of FBXL5 with VHL promotes VHL poly-ubiquitination and degradation, events that depend on the assembly of FBXL5 into an SCF complex. The interaction itself is mediated by the C-terminus of FBXL5 (Fig. S3D) and a region spanning amino acids 114-170 of VHL with VHL mutants lacking this region being resistant to FBXL5 activity. Importantly, the FBXL5-dependent degradation of VHL profoundly impairs the ability of VHL to catalyze the ubiquitin-dependent degradation of HIF1α leading to HIF1α stabilization and upregulation of the hypoxia gene expression program. Taken together, these observations identify FBXL5-VHL as novel regulatory axis that functions redundantly with the canonical PHD-mediated regulation of VHL-HIF1α, to tightly regulate the cellular response to hypoxia.

**Figure 6.**
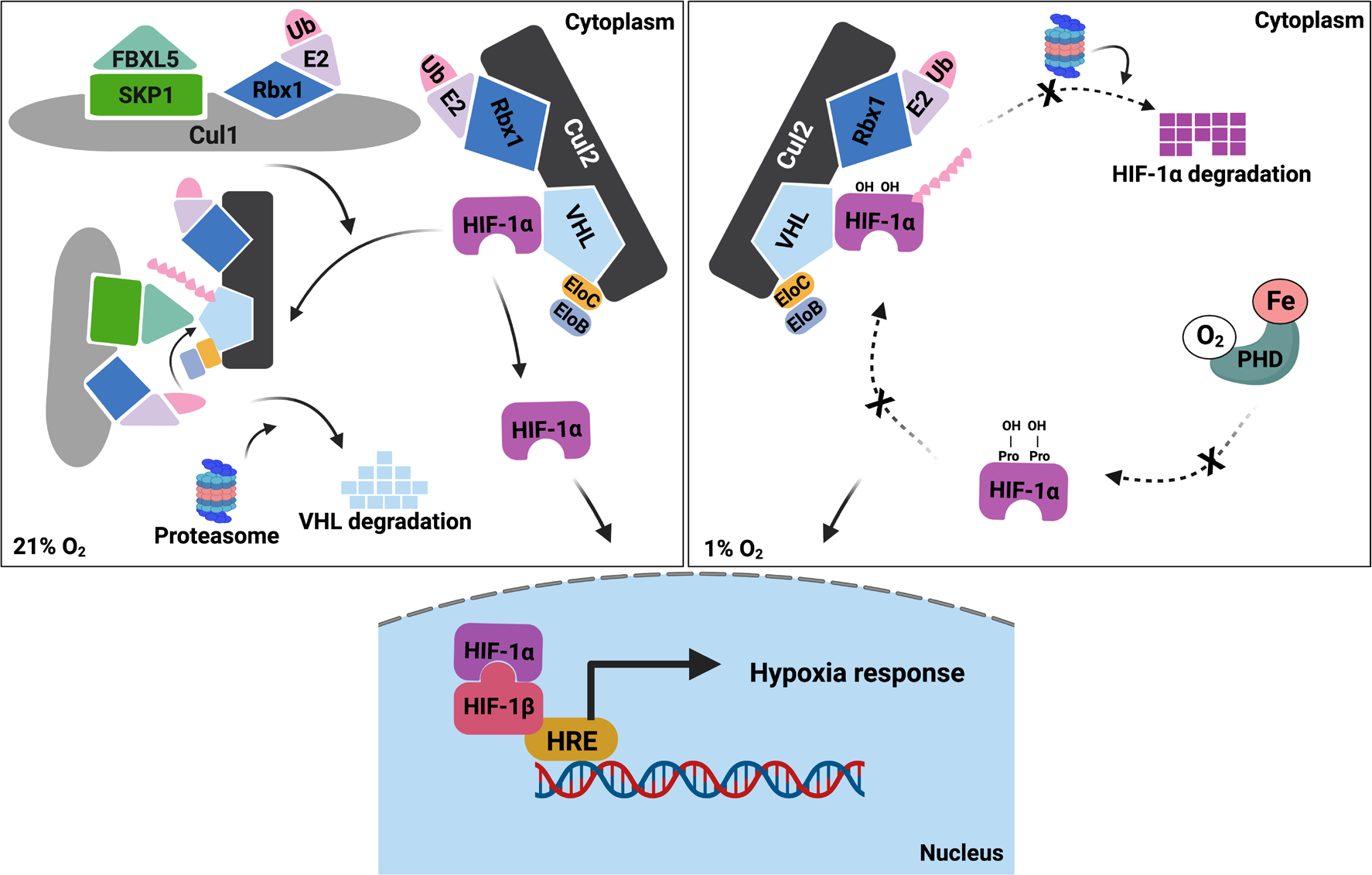
FBXL5-mediated VHL regulation: A redundant pathway to HIF1α stabilization. SCF^FBXL5^ complex interacts with VHL protein via substrate adaptor FBXL5 and promotes its ubiquitin-dependent degradation leading to HIF1α stabilization. The existence of parallel pathways for inactivation of PHDs in hypoxia and VHL degradation by FBXL5 ensures rapid accumulation of HIF1α, which in turn promotes transcriptional activation of genes involved in the cellular adaptive hypoxia response.

Prolyl-hydroxylases catalyze the hydroxylation of HIFs in a reaction that depends on both molecular O_2_ and iron (Bruick and McKnight, 2001; Fong and Takeda, 2008). As such, hypoxia or iron starvation act as environmental cues that directly inhibit PHDs leading to the impaired hydroxylation and reduced degradation by VHL. Although PHDs play a major role in linking intracellular iron and oxygen levels to HIFα protein stability, our studies identify a second regulatory axis mediated by FBXL5 and VHL that also responds to changes in iron and oxygen levels and cooperates with PHDs to regulate HIF stability. Importantly, these regulatory events appear to function largely independent of each other. For example, cells depleted of FBXL5 showed downregulation of HIF1α, even in the presence of PHD2 inhibitor dimethyl-oxalylglycine (DMOG) (Fig. S6A). Additionally, the up-regulation of HIF1α levels in cells overexpressing wild-type FBXL5 was unaffected by siRNA-mediated knockdown of PHD2 (Fig. S6B). On the contrary, this effect was dampened by overexpression of the dominant negative FBXL5-ΔFbox mutant which blocks VHL degradation. Further, we did not observe significant effects of either wild-type or FBXL5-ΔFbox mutant overexpression on PHD2 levels (Fig. S6B). These data suggest that HIF1α accumulation is dependent on both PHDs to regulate HIF1α hydroxylation and FBXL5 to regulate VHL abundance. The independent nature of these regulatory events is consistent with a model in which HIF stabilization is tightly controlled by multiple redundant mechanisms that ensure HIF1α is not inappropriately activated outside of hypoxia and iron starvation conditions. Further investigation is needed to tease out the non-redundant roles of FBXL5 and PHDs to better understand their individual contributions to the regulation of HIF1α-dependent hypoxia response.

Cellular levels of molecular oxygen and iron have long been shown to cross-regulate each other with significant consequences for many biological processes. One clear avenue of cross regulation is via the IRP-IRE system. For example, many genes including HIF2α that are tasked with modulating the cellular response to oxygen bioavailability contain IREs in their 5’ or 3’ untranslated regions (UTR) and are regulated post-transcriptionally by IRP binding (Sanchez et al., 2007; Shah et al., 2009). In this context, it is clear in our study that FBXL5 can impact oxygen homeostasis through distinct mechanisms that include regulation of the IRE-IRP axis through IRP degradation and the VHL-HIF axis through VHL degradation. Despite the challenges associated with separating these regulatory arms, the data clearly support the independence of these pathways. First, HIF1α has not been reported to possess an IRE and its mRNA levels are not responsive to changes in the FBXL5-IRP axis (Fig. S4B) supporting that its regulation by FBXL5 reported here occurs independently of IRPs. Second, we have generated mutants of FBXL5 that have lost the ability to degrade IRPs and shown that these mutants are still able to degrade VHL (Fig 5E and Fig. S5E) and control HIF1α abundance (Fig 5C) consistent with these functions being separable. Finally, siRNA-mediated silencing of IRP2 in cells overexpressing either wild-type or FBXL5-ΔFbox mutant under hypoxia conditions failed to display any significant effects on either HIF1α or VHL levels compared to controls (Fig. S7). Together, these data support a model in which the ability of FBXL5 to influence HIF1α stability via VHL degradation is independent of the IRP-IRE system at least with respect to HIF1α. Further work will be required to assess the relative contribution of these two regulatory arms on influencing levels of HIF2α which possesses an IRE and whose stability is clearly affected by FBXL5 (Fig. S8).

Although HIF1α protein stability has been the long-standing focus of hypoxia response field, understanding the regulation of adaptor protein VHL is crucial to addressing the current gaps in our knowledge of hypoxia-related pathologies. Multiple studies have previously reported mechanisms for regulating VHL by protein degradation. GAM1, an adenoviral substrate receptor, was shown to promote degradation of VHL protein in a ubiquitin- and proteasome-dependent manner (Pozzebon et al., 2013). Tumor-disposing mutants of VHL defective in elongin binding have been showed to destabilize VHL leading to the impaired degradation of HIF substrates (Kamura et al., 2002; Schoenfeld et al., 2000). Of note, Kamura et al. showed that Rbx1 interacted with VHL in the absence of exogenously expressed Cul2, Elongins B and C in Sf21 insect cells and that VHL ubiquitylation was dependent on functional Rbx1. This observation can be potentially explained by our finding that VHL interacts with SCF^FBXL5^ which in turn associates with ubiquitin conjugating E2 enzyme via Rbx1. To achieve effective ubiquitination of VHL, the F-Box domain of FBXL5 is crucial for mediating its assembly into SKP1-Cul1-Rbx1 complex. Impairment of this assembly by either deletion of F-box domain (this study) or a mutant Rbx1 incompetent in recruiting E2 enzyme (Kamura et al.) substantially reduces ubiquitination of VHL.

## Acknowledgements

This work was supported by the National Institutes of Health GM089778 to JAW.

## Author contributions

Conceptualization, A.K.M, V.P., J.A.W; Methodology, A.K.M., V.P., J.A.W.; Investigation, A.K.M, V.P., K.L., L.L., Y.J., A.M.K.; Writing, V.P., A.K.M, J.A.W; Funding Acquisition, J.A.W.

## Star Methods

Further information and requests for resources and reagents should be directed to and will be fulfilled by the Lead Contact, James A. Wohlschlegel.

## Experimental Model and Subject Details

HEK293 (ATCC, sex: female), Flp-In™ T-REx™ 293 cells (ThermoFisher Scientific, sex: female) and Flp-In™ T-REx™ Hela cells (ThermoFisher Scientific, sex: female) were utilized for the study. All cell lines were maintained at 37°C, 5% CO_2_ in DMEM containing 4.5g glucose/L (Gibco by Life Technologies) supplemented with 10% fetal bovine serum (Gemini Bioproducts), 1X antibiotic-antimycotic (Gibco by Life Technologies) and 2mM L-glutamine (Gibco by Life Technologies). For hypoxia studies, cells were maintained in a gaseous mixture of 1% O_2_, 5% CO_2_ and 94% N_2_ at 37°C for 16 hours. To induce gene expression in Flp-In™ T-REx™ 293 cells, doxycycline (Fisher Scientific) was added to the culture media at a final concentration of 1mg/mL every 24 hours before harvesting cells for analysis as indicated.

## Method Details

### Plasmids

The generation of pcDNA5-FRT/TO-3xHA-3xFLAG-FBXL5, pcDNA3-6xMyc-FBXL5 and pcDNA3-6xMyc-FBXL5-ΔFbox has been described earlier (Stehling et al., 2012; Vashisht et al., 2009). pDONR221-FBXL5 (described previously in (Vashisht et al., 2009) was subcloned into pEYFP-N1 DEST plasmid to generate pEYFP-N1-FBXL5 using Gateway cloning system (Invitrogen). The human FBLX5 ORF with deletions from 666-691aa, 651-691aa (ΔLRR7), 623-691aa (ΔLRR6,LRR7) and cysteine to serine mutations at C662, C676, C686 and C687 (as described in (Wang et al., 2020)) were ordered from Integrated DNA Technologies (IDT) and cloned into pDONR221 followed by subcloning into DEST plasmid pcDNA3-6xMyc to generate pcDNA3-6xMyc-FBXL5-Δ666-691, pcDNA3-6xMyc-FBXL5-ΔLRR7, pcDNA3-6xMyc-FBXL5-ΔLRR6,LRR7 and pcDNA3-6xMyc-FBXL5-4C>S. 2xFLAG-IRP2 and HA-Ub were previously described (Johnson and Blobel, 1997; Zumbrennen et al., 2009). Using 2xFLAG-IRP2 as a template, pcDNA3-6xMyc-IRP2 was generated using Gateway cloning system (Invitrogen). pDONR223-VHL (a gift from Jesse Boehm & William Hahn & David Root, Addgene plasmid # 81874)(Kim et al., 2016) was subcloned using Gateway system (Invitrogen) to generate pcDNA3-6xMyc-VHL, pEYFP-VHL and pgLAP3-EGFP-VHL (Torres et al., 2009). ORFs of various deletion mutants of Human VHL flanked by attB1 and B2 sequences as shown in Table S1 were synthesized either by IDT or Twist Biosciences, cloned into pDONR221, and further subcloned into either pgLAP3-EGFP-N1 or pEYFP-N1 using Gateway Technology. For VHL-K3R and K196R mutants, ORFs were synthesized from Twist Biosciences incorporating lysine to arginine mutations at amino acids at K159, K171 and K196 or at K196 alone flanked by attB1 and B2 sequences were cloned into pDONR221 and pEYFP-N1 using Gateway Technology.

### Generation of stable cell lines

Generation of Flp-In™ T-REx™ 293 cells stably expressing 3xHA-3xFLAG-FBXL5, 3xHA-3xFLAG-FBXL5-ΔFbox, 3xHA-3xFLAG-FBXL5-C500, 3xHA-3xFLAG-FBXL5-ΔCTBD and 3xHA-3xFLAG-N199 have been described previously (Mayank et al., 2019; Stehling et al., 2012; Vashisht et al., 2009). Flp-In™ T-REx™ 293 cells stably expressing 3xHA-3xFLAG-VHL and 3xHA-3xFLAG-FBXL5-4C>S as well as Flp-In™ T-Rex™ Hela cells stably expressing 3xHA-3xFLAG-FBXL5 and 3xHA-3xFLAG-FBXL5-ΔFbox were generated using the Flp-In system (Invitrogen) according to the manufacturer’s protocol. Generation of *FBXL5*-KD and *FBXL5*-KD^FBXL5-WT^ CRISPRi cell lines have been described previously (Mayank et al., 2019).

### Affinity purification of FBXL5 and VHL-containing protein complexes

Six 15 cm tissue culture plates each of Flp-In 293 or Flp-In 293 stably expressing 3XHA-3-Flag-tagged FBXL5, ΔFbox and VHL were grown, harvested and lysed in IP buffer composed of 100mM Tris-HCl pH8.0, 150mM NaCl, 50mM EDTA, 0.1% NP-40, 10% glycerol, 1mM DTT, protease inhibitors and phosphatase inhibitors. Clarified protein lysates were normalized by protein concentration and then incubated with 100uL of EZ-view anti-HA agarose (Pierce) for 2 hours at 4°C. Beads were washed four times using 1 mL of IP buffer per wash followed by washes with IP buffer lacking NP-40. Following the washes, bound protein complexes were eluted using 8M urea and 100mM Tris-Cl pH 8.0 and then precipitated in acetone.

### Proteomic characterization of affinity purified FBXL5 samples

Acetone precipitates were resuspended in 100mM Tris-HCl pH 8.5, 8M urea, reduced and alkylated using 5mM Tris (2-carboxyethyl) phosphine and 10mM iodoacetamide, respectively, and proteolytically digested with lys-C and trypsin. The digested peptides were subsequently analyzed by LC-MS/MS. Briefly, peptides were separated by reversed phase chromatography using Bruker PepSep C18 Columns (150μm ID, 15 cm lengths, 1.5 μm particle size) using a gradient of increasing acetonitrile delivered by a Vanquish Neo UHPLC-System (Thermo Scientific) operating at a flow rate of 500nL/min. Data was acquired on a Bruker timsTOF HT mass spectrometer using a diaPASEF data acquisition strategy. The dia-PASEF windows spanned from 300 m/z to 1200 m/z using the following ion mobility parameters: 1/K0 start: 0.60 Vs cm−2; 1/K0 end: 1.60 Vs cm−2; ramp time: 100 ms; accumulation time: 50 ms. The dia-PASEF files were searched with DIA-NN using an *in silico* library generated from a Uniprot database containing all human proteins (Demichev et al., 2020). DIA-NN output files were subsequently analyzed by the MSStats R package to identify differentially enriched proteins (Choi et al., 2014). The mass spectrometry data have been deposited to MassIVE repository with the identifier MSV000093946.

### Immunoprecipitation

Cell lysates were prepared using immunoprecipitation (IP) buffer consisting of 100mM Tris-HCl pH 8.0, 150mM NaCl, 50mM EDTA, 0.1% NP-40, 10% glycerol, 1mM DTT, protease inhibitors and phosphatase inhibitors. Equal protein amounts were incubated for 2 hours at 4°C with appropriate affinity matrix pre-equilibrated using IP buffer. Beads were washed 4 times in IP buffer followed by eluting bound protein complexes in 2X SDS dye. For immunoblotting, immunoprecipitates and whole cell lysates were boiled in SDS loading buffer, resolved on SDS-PAGE, followed by transfer onto PVDF membrane. The membranes were blocked in either 5% milk or BSA followed by incubation with appropriate primary antibody and HRP-labeled secondary antibody. Protein bands on membranes were visualized using Pierce ECL western blotting substrate (ThermoFisher).

### Ubiquitination assay

HEK293 cells were transfected with plasmids expressing HA-Ubiquitin, VHL constructs (GFP-VHL, YFP-VHL, YFP-VHL-Δ98-170, YFP-VHL-Δ106-162, YFP-VHL-Δ114-170, YFP-VHL-Δ134-170, YFP-VHL-K3R or YFP-VHL-K196R) and either 6xMyc-FBXL5, 6xMyc-FBXL5-ΔFbox, 6xMyc-FBXL5-4C>S or vector control. Twenty-four hours post-transfection cells were treated with 10 μM MG132 for 2 hours or as indicated. Cells were harvested and lysed under denaturing conditions as described previously (Bloom and Pagano, 2005). Ubiquitin conjugates were purified using anti-HA beads and the presence of VHL and its mutants in the purified ubiquitin conjugates was detected by immunoblotting with GFP antibody.

### Quantification of mRNA by one-step RT-qPCR

Total RNA from cells was extracted using the Aurum™ Total RNA Mini Kit (Bio-Rad, CA, USA), and 100ng of RNA was used with iTaq™ Universal SYBR® Green One-Step Kit (Bio-Rad, CA, USA) to perform a one-step RT-qPCR in a volume of 20 μl using a QuantStudio3 real-time PCR instrument from ThermoFisher (Waltham, MA, USA). The primers used are provided in the Key Resource Table. β-actin was used to normalize the Ct values, which were then used to calculate fold changes using the ΔΔCt method.

### Cell transfections and treatments

HEK293 cells were transiently transfected with indicated plasmids using either BioT (Bioland, Long Beach, CA), Lipofectamine™2000 or Lipofectamine™3000 according to the manufacturers’ protocol. Cells were treated with the following drugs/inhibitors at the indicated concentrations and for the time periods listed (unless indicated otherwise in the results section): FAC (100μg/mL) for 8 hours; DFO (100μM) for 8 hours, MG132 (10μM) for 2 hours, DMOG (0.5mM) for 24 hours, CHX (100μg/mL) at indicated time-points, BafA (10nM) for 8 hours and CoCl_2_ (100μM) for 16 hours. siRNA transfections were carried out using Lipofectamine®RNAiMax as the transfection reagent and siGENOME SMARTpool FBXL5 siRNA (Horizon #M-012424-01-0005), siGENOME SMARTpool VHL siRNA (Horizon M-003936-00-0005), siGENOME SMARTpool PHD2 siRNA (Horizon L-004276-00-0005), siGENOME SMARTpool IREB2/IRP2 siRNA (Horizon M-022281-01-0005), and siGENOME non-targeting siRNA (Horizon #D-001210-03-20) based on the manufacturer’s instructions, For hypoxia treatment, cells growing at 60% confluence were transferred to a hypoxia chamber (STEMCELL technologies) which was purged with a gaseous mixture of 1% O_2_, 5% CO_2_ and 94% N_2_ for 7 minutes at a rate of 20 L/min using Single Flow Meter (STEMCELL Technologies, Cat. #27311). The sealed chamber was then transferred to 37°C and cells incubated for 16 hours before harvesting for further analysis.

