## Supplemental Data for "SCF^FBXL5^ ubiquitin ligase regulates stability of von-Hippel Lindau protein and the HIF1α-dependent response to hypoxia"

### Supplemental Figures

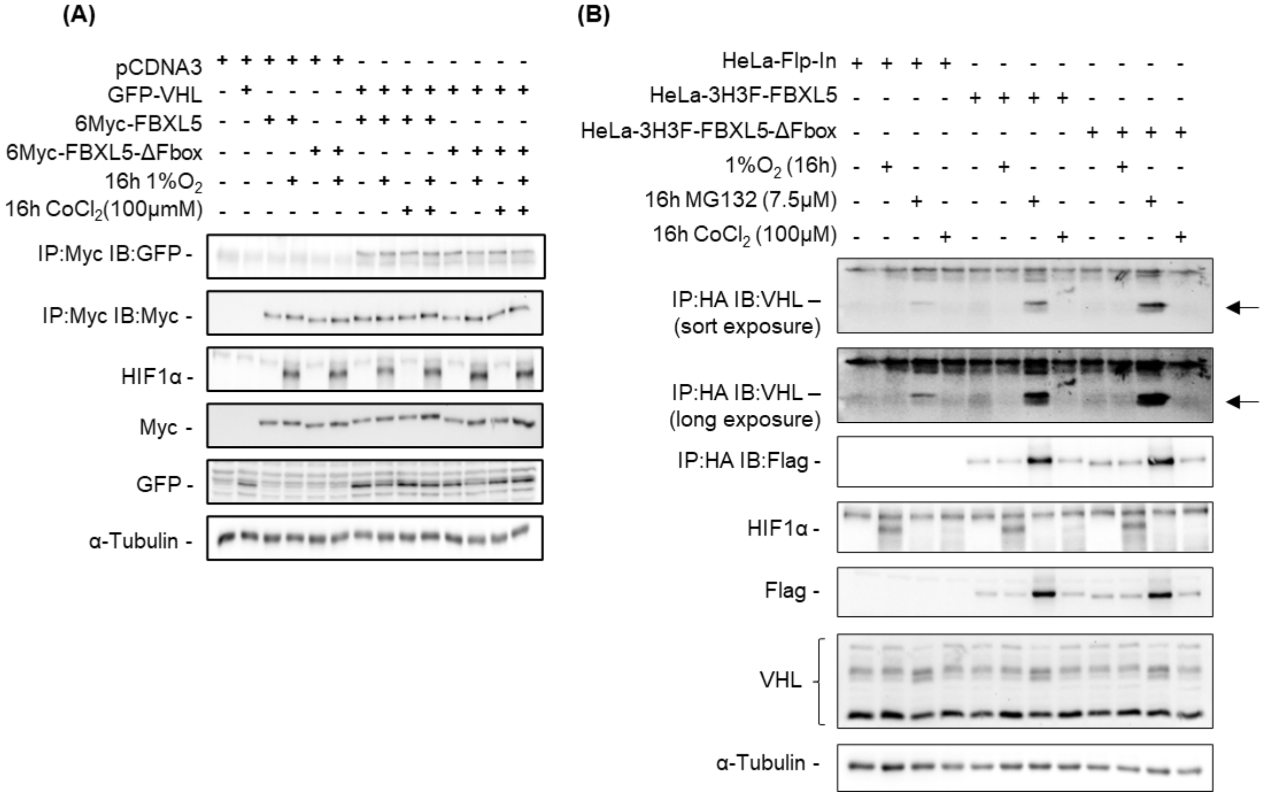


**Figure S1. Interaction between FBXL5 and VHL in Hela cells (related to Figure 1).** (A) HeLa-Flp-In cells were co-transfected with GFP-VHL and either Myc-FBXL5 or Myc-FBXL5-ΔFbox. 24 hours post-transfections, cells were either treated with 100μM $CoCl_2$ or exposed to 1% $O_2$ or combination of both for 16 hours before harvesting. Whole cell extracts (WCEs) were immunoprecipitated using beads conjugated to anti-Myc antibodies. IPs and WCEs were immunoblotted with GFP, Myc, HIF1α and α-tubulin antibodies. (B) HeLa-Flp-In cells stably overexpressing either HA-Flag-FBXL5 or HA-Flag-FBXL5-ΔFBox were treated with either 100μM $CoCl_2$ and 7.5μM MG132 or exposed to 1% $O_2$ for 16 hours. Whole cell extracts were immunoprecipitated using beads conjugated to anti-HA antibodies. IPs and WCEs were immunoblotted with VHL, Flag, HIF1α and α-tubulin antibodies.

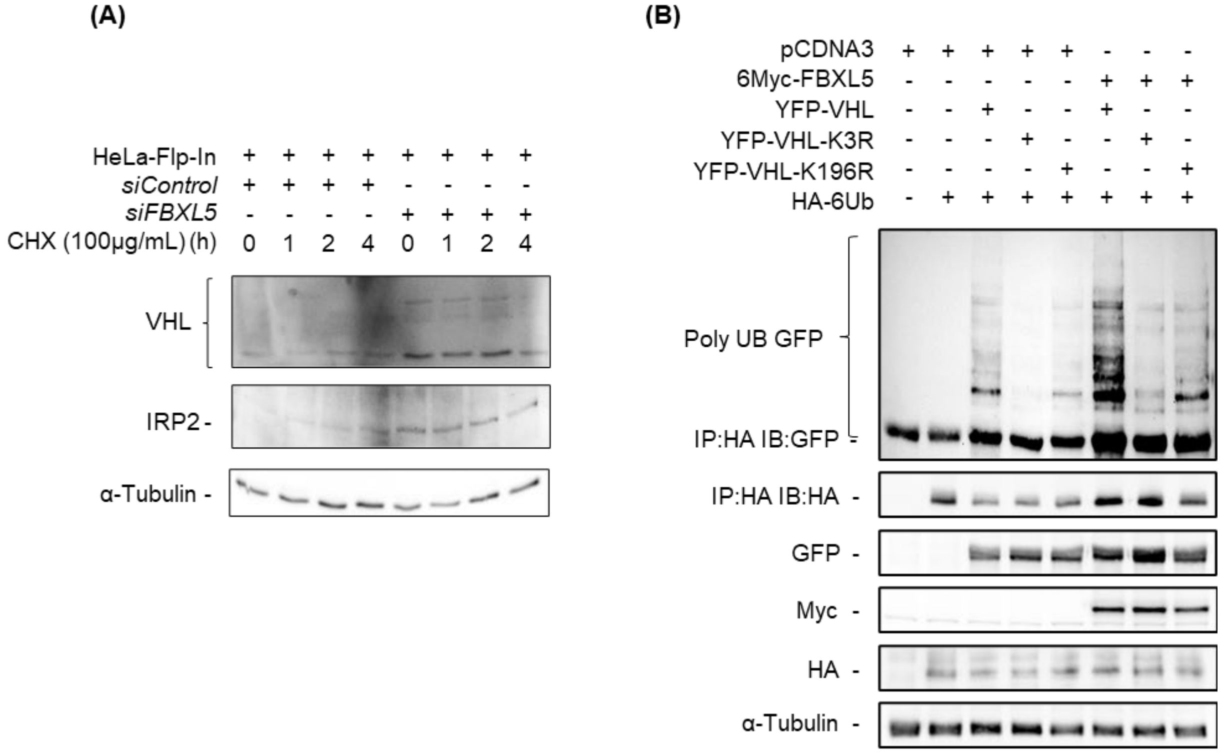


**Figure S2. FBXL5 negatively affects VHL stability (related to Figure 2).** (A) HeLa Flp-In cells transfected with either control siRNA or siRNA against *FBXL5* were treated with 100ug/mL of cycloheximide (CHX). Post-CHX treatment, cells were harvested at indicated time-points. Whole cell extracts were immunoblotted with VHL, IRP2 and α-Tubulin antibodies. IRP2 stabilization was used to indicate the efficacy of siRNA-mediated depletion of FBXL5. (B) HEK293 cells were co-transfected with HA-Ub and Myc-FBXL5 along with either YFP-VHL-WT, YFP-VHL-K3R or YFP-VHL-K196R. Post-transfection, cells were treated with 15µM MG132 for 4 hours before harvesting. Ubiquitin conjugates were immunoprecipitated by using anti-HA antibodies. HA immunoprecipitates (IP: HA) and WCEs were immunoblotted with GFP, HA, Myc and α-tubulin antibodies.

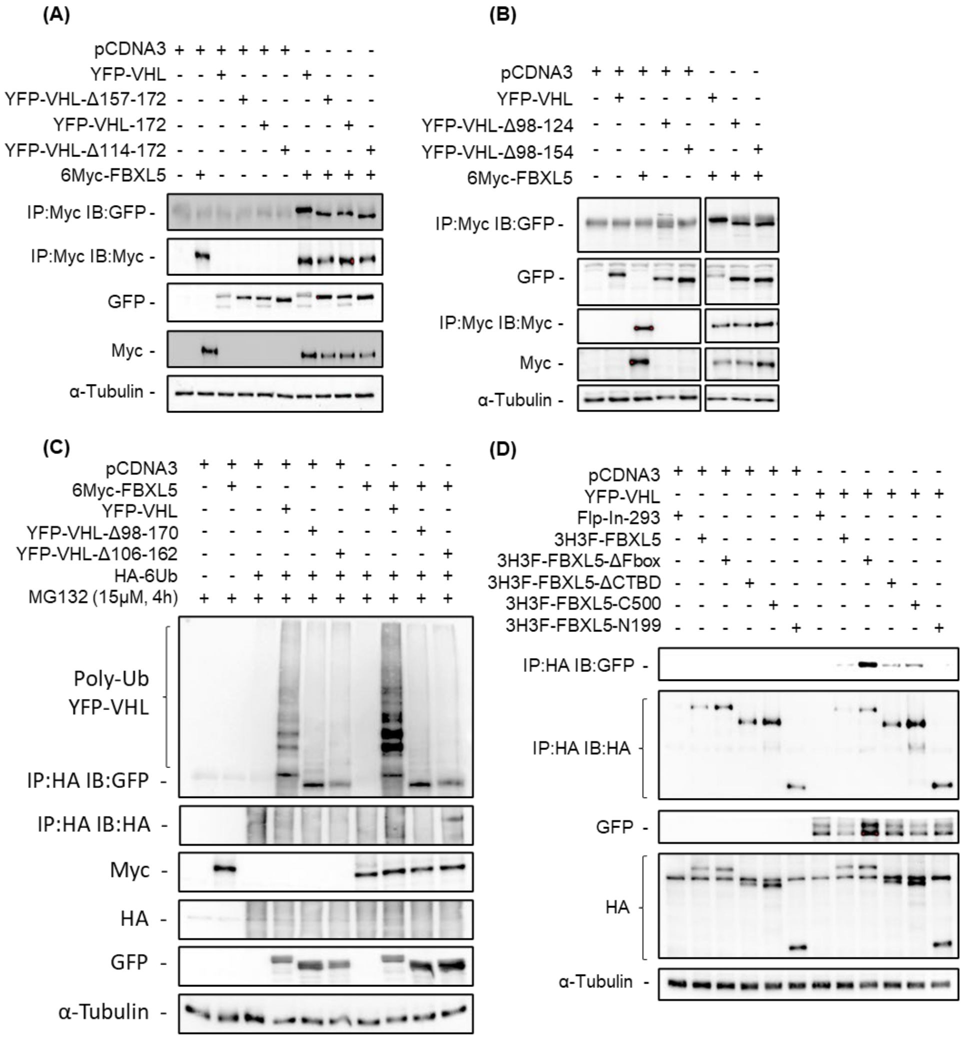


**Figure S3. FBXL5-VHL interaction site mapping (related to Figure 3).** (A) HEK293 cells were co-transfected with Myc-FBXL5 and either YFP-VHL-WT, YFP-VHL-172, YFP-VHL-Δ114-172 or YFP-VHL-Δ157-172. Whole cell extracts (WCEs) were immunoprecipitated using beads conjugated to anti-Myc antibodies. Myc-immunoprecipitates (IP:Myc) and WCEs were immunoblotted using antibodies against Myc, GFP and α-tubulin. (B) HEK293 cells were co-transfected with Myc-FBXL5 and either YFP-VHL-WT, YFP-VHL-Δ98-124 or YFP-VHL-Δ98-154. WCEs were immunoprecipitated using beads conjugated to anti-Myc antibodies. Myc-immunoprecipitates (IP:Myc) and WCEs were immunoblotted using antibodies against Myc, GFP and α-tubulin. (C) HEK293 cells were co-transfected with HA-Ub and Myc-FBXL5 along with either YFP-VHL-WT, YFP-VHL-WT, YFP-VHL-Δ98-170 or YFP-VHL-Δ98-162. Post-transfection, cells were treated with 15μM MG132 for 4 hours before harvesting. Ubiquitin conjugates were immunoprecipitated by using anti-HA antibodies. HA immunoprecipitates (IP: HA) and WCEs were immunoblotted with GFP, HA, Myc and α-tubulin antibodies. (D) HEK-Flp-In-293 cells stably overexpressing either HA-Flag-FBXL5, HA-Flag-FBXL5-ΔFbox, HA-Flag-FBXL5-ΔCTBD, HA-Flag-FBXL5-C500 or HA-Flag-FBXL5-N199 were transfected with either pCDNA3 or YFP-VHL. Whole cell lysates were immunoprecipitated using beads conjugated to anti-HA antibodies. HA-immunoprecipitates (IP: HA) and WCEs were immunoblotted using antibodies against HA, GFP and α-tubulin.

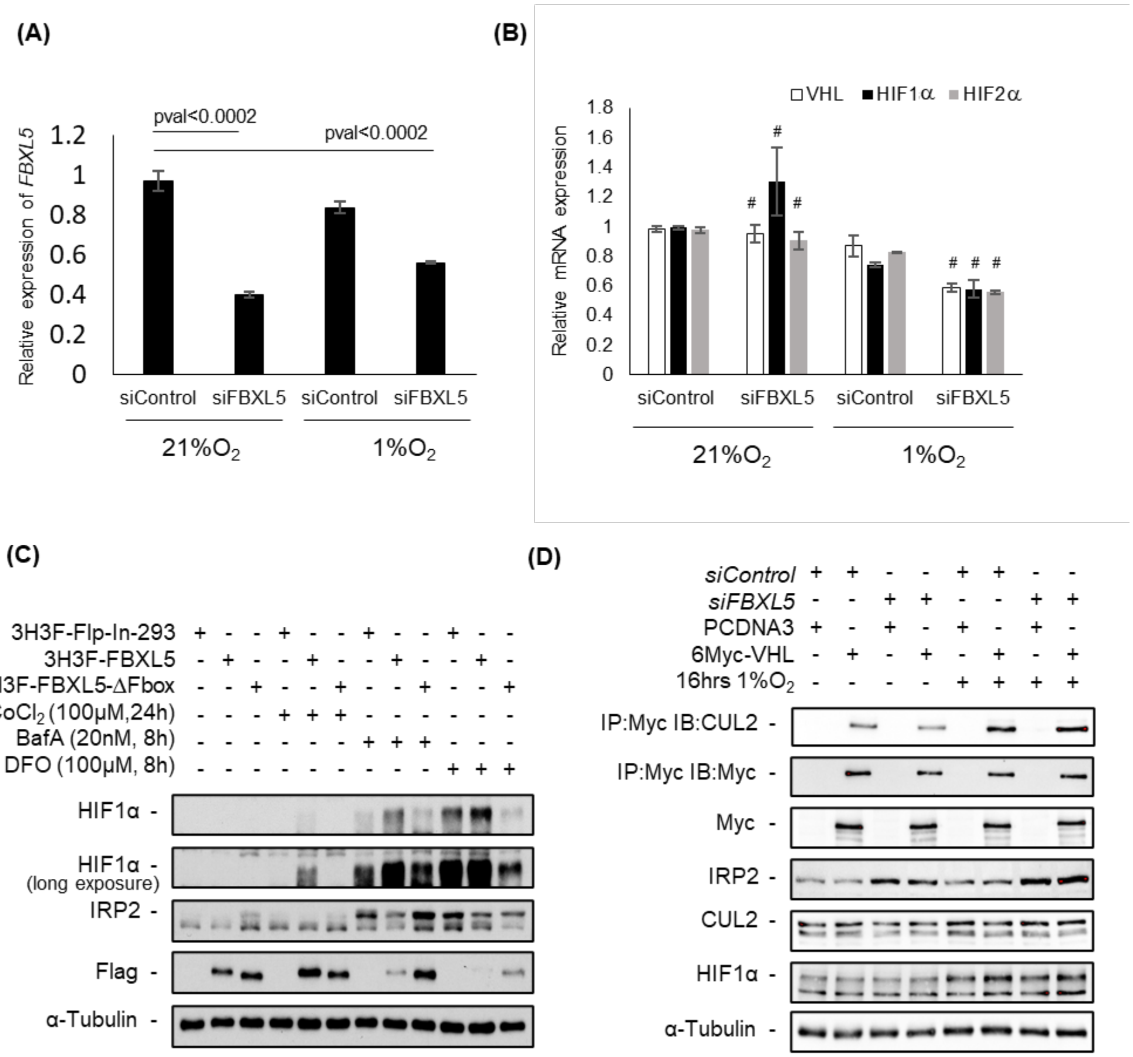


**Figure S4. Effect of FBXL5 knockdown on HIF1α protein levels (related to Figure 4).** (A) HEK293 cells were treated with either control siRNA or siRNA against *FBXL5*. Post-transfection, cells were exposed to either 21% or 1% $O_2$ for 16 hours. Relative mRNA levels of FBXL5 were determined using qPCR. (B) HEK293 cells were treated with either control siRNA or siRNA against *FBXL5*. Post-transfection, cells were exposed to either 21% or 1% $O_2$ for 16 hours. Relative mRNA levels of FBXL5 were determined using qPCR. $^{\#}$denotes p-value > 0.05 when compared to control siRNA. (C) HEK-Flp-In-293 cells stably overexpressing either HA-Flag-FBXL5 or HA-Flag-FBXL5-ΔFbox were treated with $CoCL_2$, BafA or DFO with concentrations and time as indicated. WCEs were immunoblotted using antibodies against HIF1α, IRP2 and α-tubulin. (D) HEK293 cells were treated with either control siRNA or siRNA against *FBXL5* followed by co-transfection with Myc-VHL. Post-transfection, cells were exposed to either 21% or 1% $O_2$ for 16 hours before harvesting. Whole cell extracts were immunoprecipitated using beads conjugated to anti-Myc antibodies. IPs and WCEs were immunoblotted with CUL2, Myc, HIF1α, IRP2 and α-tubulin antibodies. IRP2stabilization was used to indicate the efficacy of siRNA-mediated depletion of FBXL5.

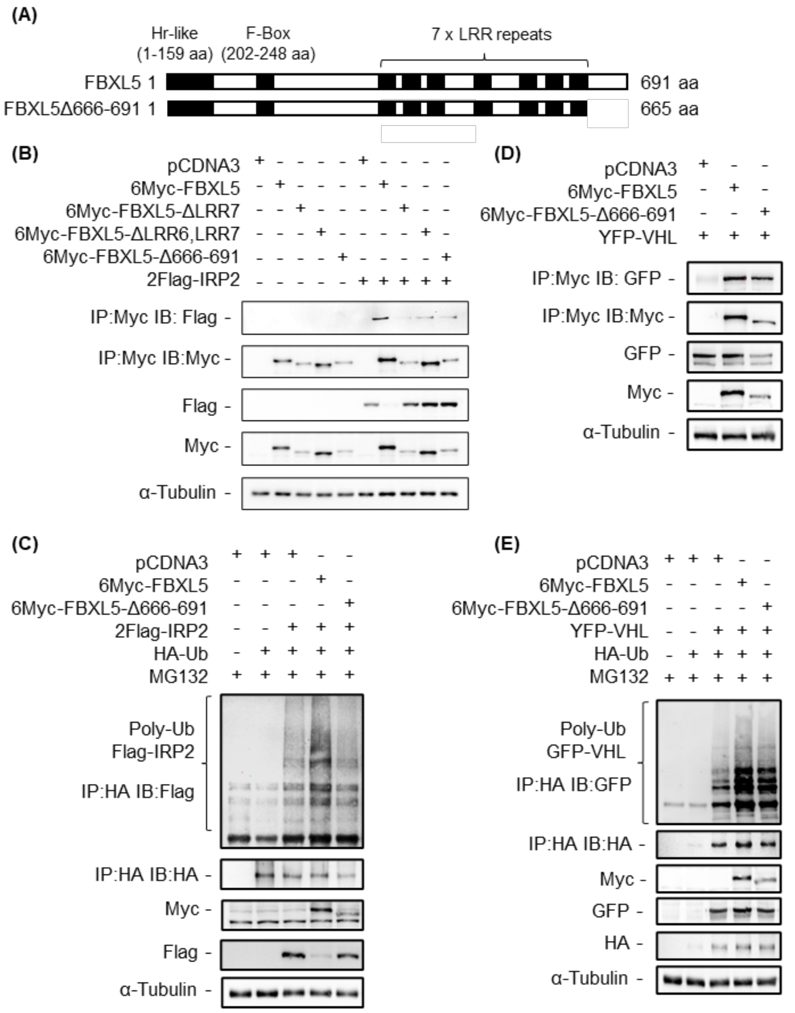


**Figure S5. SCF$^{FBXL5}$ complex employs distinct mechanisms for regulation of VHL and IRP2 protein stability (related to Figure 5).** (A) Domain structure of FBXL5 C-terminal deletion mutant. (B) HEK293 cells were co-transfected with Flag-IRP2 and either Myc-FBXL5, Myc-FBXL5-ΔLRR7, Myc-FBXL5-ΔLRR6&7 or Myc-FBXL5-Δ666-691. Whole cell extracts (WCEs) were immunoprecipitated using beads conjugated to anti-Myc antibodies. Myc-immunoprecipitates (IP:Myc) and WCEs were immunoblotted using antibodies against Myc, Flag and α-tubulin. (C) HEK293 cells were co-transfected with HA-Ub and Flag-IRP2 along with either Myc-FBXL5 or Myc-FBXL5-Δ666-691. Post-transfection, cells were treated with 15μM MG132 for 4 hours before harvesting. Ubiquitin conjugates were immunoprecipitated by using anti-HA antibodies. HA immunoprecipitates (IP: HA) and WCEs were immunoblotted with Flag, HA, Myc and α-tubulin antibodies. (D) HEK293 cells were co-transfected with YFP-VHL and either Myc-FBXL5 or Myc-FBXL5-Δ666-691. WCEs were immunoprecipitated using beads conjugated to anti-Myc antibodies. Myc-immunoprecipitates (IP:Myc) and WCEs were immunoblotted using antibodies against Myc, GFP and α-tubulin. (E) HEK293 cells were co-transfected with HA-Ub and YFP-VHL along with either Myc-FBXL5 or Myc-FBXL5-Δ666-691. Post-transfection, cells were treated with 15μM MG132 for 4 hours before harvesting. Ubiquitin conjugates were immunoprecipitated using anti-HA antibodies. HA immunoprecipitates (IP: HA) and WCEs were immunoblotted with GFP, HA, Myc and α-tubulin antibodies.

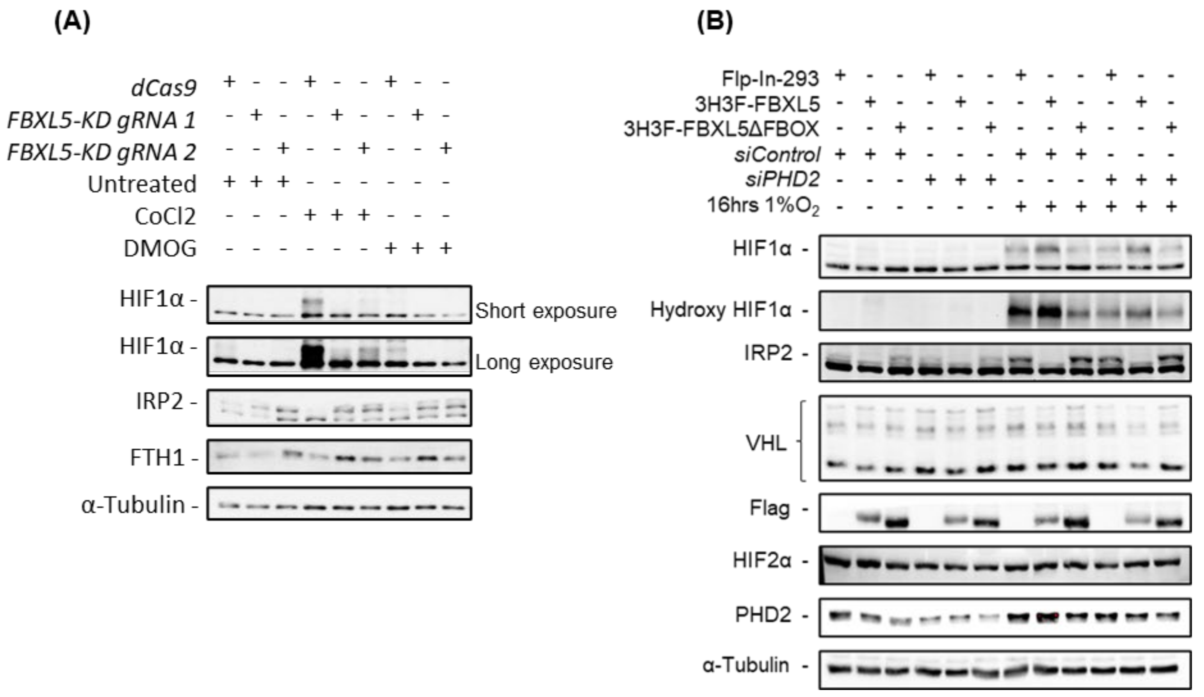


**Figure S6. Effect of PHD2 knockdown and PHD inhibitor on HIF1α expression in FBXL5 over-expressing cells under hypoxic conditions (related to discussion).** (A) HEK-Flp-In-293 cells stably overexpressing either HA-Flag-FBXL5 or HA-Flag-FBXL5-ΔFbox were transfected with either control siRNA or siRNA against *PHD2*. 24 hours post-transfections, cells were exposed to 1% $O_2$ for 16 hours. WCEs were immunoblotted using antibodies against VHL, Flag, HIF1α, HIF2α, PHD2, IRP2 and α-tubulin. IRP2 stabilization was used to indicate the efficacy of siRNA-mediated depletion of FBXL5. (B) Flp-In 293 cells were infected with lentiviral vectors expressing dCas9 with non-target (dCas9) or FBXL5 specific gRNA 1 and 2 (*FBXL5-KD gRNA1 and FBXL5-KD gRNA2*). Cells were treated with $CoCL_2$ (100μM, 16 hours) and DMOG (0.5mM, 24 hours). WCEs were immunoblotted with HIF1α, IRP2 and α-tubulin antibodies.

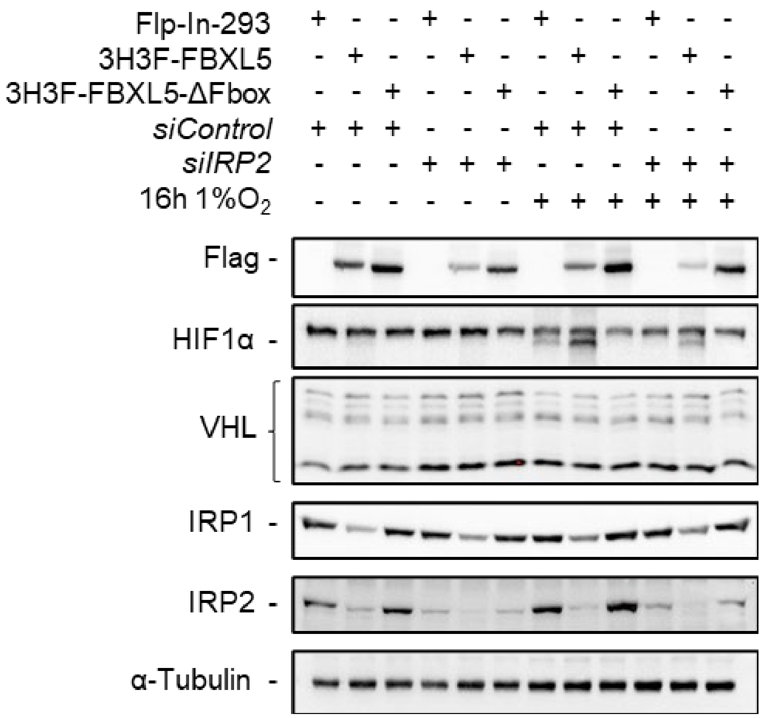


**Figure S7. Effect of IRP2 knockdown on HIF1α expression in FBXL5 over-expressing cells under hypoxic conditions (related to discussion).** (A) HEK-Flp-In-293 cells stably overexpressing either HA-Flag-FBXL5 or HA-Flag-FBXL5-ΔFbox were transfected with either control siRNA or siRNA against *IRP2*. 24 hours post-transfections cells were exposed to 1% $O_2$ for 16 hours. WCEs were immunoblotted using antibodies against VHL, Flag, HIF1α, IRP1, IRP2 and α-tubulin.

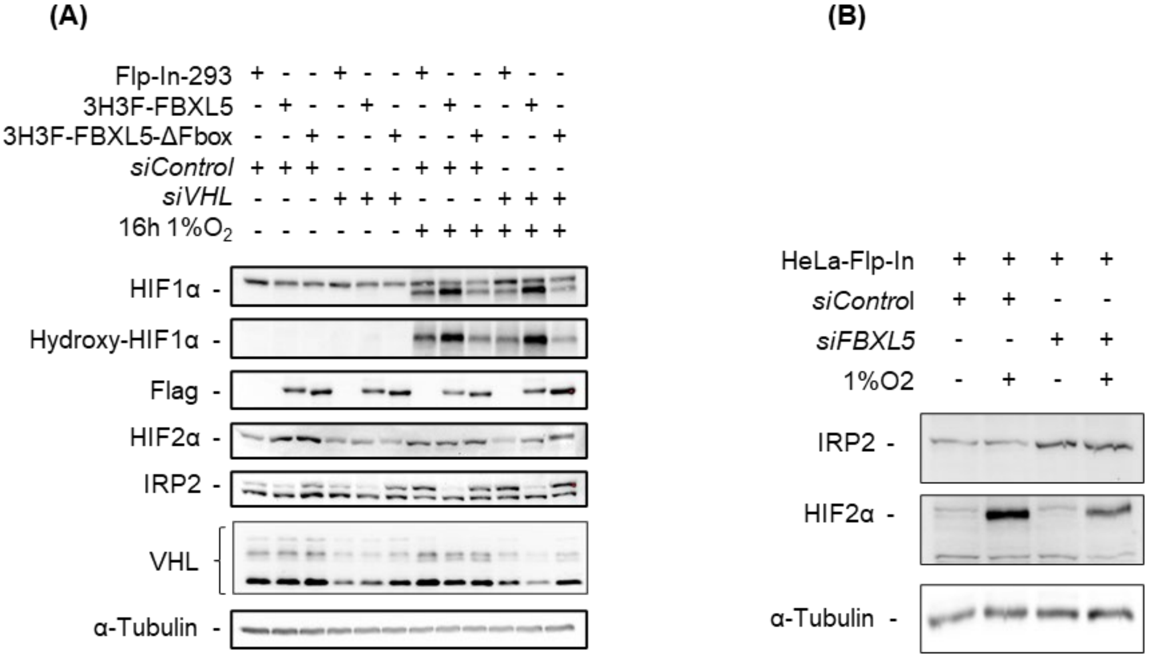


**Figure S8. Effect of VHL and FBXL5 knockdown on HIF2α expression in hypoxic conditions (related to discussion).** (A) HEK-Flp-In-293 cells stably overexpressing either HA-Flag-FBXL5 or HA-Flag-FBXL5-ΔFbox were transfected with either control siRNA or siRNA against VHL. 24 hours post-transfections, cells were exposed to 1% $O_2$ for 16 hours. WCEs were immunoblotted using antibodies against VHL, Flag, HIF1α, Hydroxy-HIF1α, HIF2α, PHD2, IRP2 and α-tubulin. (B) HeLa-Flp-In cells transfected with either control siRNA or siRNA against FBXL5. 24 hours post-transfections, cells were exposed to 1% $O_2$ for 16 hours. WCEs were immunoblotted using antibodies against HIF2α, PHD2 and α-tubulin. IRP2 stabilization was used to indicate the efficacy of siRNA-mediated depletion of FBXL5.

**Table S1**

| VHL deletion mutants | DEST plasmids | Sequence (amino acids) |
|---|---|---|
| VHL-213 | pEYFP-N1 and pgLAP3-EGFP | 1-213 |
| VHL-(172) | pEYFP-N1 and pgLAP3-EGFP | 1-113, 155-213 |
| VHL-(160) | pEYFP-N1 and pgLAP3-EGFP | 54-213 |
| VHL-114-154 | pEYFP-N1 and pgLAP3-EGFP | 114-154 |
| VHL-1-53 | pEYFP-N1 and pgLAP3-EGFP | 1-53 |
| VHL-1-113 | pEYFP-N1 and pgLAP3-EGFP | 1-113 |
| VHL-155-189 | pEYFP-N1 and pgLAP3-EGFP | 155-189 |
| VHL-Δ98-124 | pEYFP-N1 and pgLAP3-EGFP | 1-97, 125-213 |
| VHL-Δ98-154 | pEYFP-N1 and pgLAP3-EGFP | 1-97, 155-213 |
| VHL-Δ98-170 | pEYFP-N1 and pgLAP3-EGFP | 1-97, 171-213 |
| VHL-Δ106-154 | pEYFP-N1 and pgLAP3-EGFP | 1-105, 155-213 |
| VHL-Δ106-162 | pEYFP-N1 and pgLAP3-EGFP | 1-105, 163-213 |
| VHL-Δ114-162 | pEYFP-N1 and pgLAP3-EGFP | 1-113, 163-213 |
| VHL-Δ114-170 | pEYFP-N1 and pgLAP3-EGFP | 1-113, 171-213 |
| VHL-(172)-Δ157-172 | pEYFP-N1 and pgLAP3-EGFP | 1-113, 155-156, 173-213 |
| VHL-Δ157-172 | pEYFP-N1 and pgLAP3-EGFP | 1-156, 173-213 |

| REAGENT or RESOURCE | SOURCE | IDENTIFIER |
|---|---|---|
| Antibodies | | |
| Monoclonal ANTI-FLAG® M2 | Sigma-Aldrich | Cat #F1804 RRID:AB_262044 |
| Rabbit monoclonal anti-Hydroxy HIF1α | Cell Signaling Tech. | Cat #(D43B5) XP® Rabbit mAb 3434 |
| Rabbit monoclonal anti-VHL | Cell Signaling Tech. | Cat #Rabbit mAb 7096 |
| Rabbit monoclonal anti-HIF2α | Cell Signaling Tech. | Cat #(D9E3) Rabbit mAb 7096 |
| Mouse monoclonal anti-IRP2 (7H6) | Santa Cruz Biotechnology | Cat #sc-33682 RRID:AB_2126703 |
| HRP-conjugated α -tubulin | Proteintech | Cat #HRP-66031 RRID:AB_2687491 |
| Mouse monoclonal anti-FBXL5 (3F5G12G9) | Biolegend | Cat #672702 RRID:AB_2564464 |
| Mouse monoclonal anti-FBXL5 (10F4H9D12) | Biolegend | Cat #672702 RRID:AB_2564464 |
| Rabbit polyclonal anti-GFP Tag | Proteintech | Cat #50430-2-AP RRID:AB_11042881 |
| Rabbit polyclonal anti-c-Myc (A-14) | Santa-Cruz Biotechnology | Cat #sc-789 RRID:AB_631274) |
| Rabbit polyclonal anti-GST tag | Proteintech | Cat #10000-0-AP RRID:AB_11042316 |
| Rabbit polyclonal anti-HIF1α | Bethyl Laboratories Inc. | Cat #A300-286A |
| Rabbit polyclonal anti-PHD2 | Bethyl Laboratories Inc. | Cat #A300-322A |
| Rabbit monoclonal anti-VHL | Cell Signaling Tech. | Cat #(E4G3M) mAb 10450 |
| Chemicals, Peptides, and Recombinant Proteins | | |
| Doxycycline Hydrochloride | Fisher Bioreagents | Cat #BP26535 |
| Hygromycin-B Gold | InvivoGen | Cat #ant-hg |
| Puromycin | InvivoGen | Cat #ant-pr |
| Cobalt (II) chloride hexahydrate | Sigma-Aldrich | CAS 7791-13-1 |
| Ammonium ferric citrate (FAC) | Fisher Bioreagents | CAS 1185-57-5 |
| Deferoxamine mesylate (DFO) | Santa-Cruz Biotechnology | CAS 138-14-7 |
| Dimethyloxalylglycine, N-(Methoxyoxoacetyl)-glycine methyl ester (DMOG) | Sigma-Aldrich | CAS 89464-63-1 |
| N-ethylmaleimide | EMD Millipore | Cat #34115 |
| MG132 | Enzo Life Sciences | Cat #BML-PI102 |
| Critical Commercial Assays | | |
| Pierce™ ECL Western Blotting Substrate | Thermo Fisher Scientific | Cat #32106 |
| SuperSignal™ West Femto Maximum Sensitivity Substrate | Thermo Fisher Scientific | Cat #34096 |
| Gateway Cloning System | Invitrogen | |
| Experimental Models: Cell Lines | | |
| HEK293 | ATCC | ATCC®CRL-1573 RRID:CVCL_0045 |
| HEK293T | ATCC | ATCC®CRL-3216 RRID:CVCL_0063 |
| Flp-In™ T-REx™ 293 | ThermoFisher | Cat #R78007 RRID:CVCL_U427 |
| Oligonucleotides | | |
| siGENOME SMARTpool FBXL5 siRNA | Horizon | #M-012424-01-0005 |
| siGENOME Non-targeting siRNA | Horizon | #D-001210-03-20 |
| ON-TARGETplus Human EGLN1 siRNA | Horizon | #L-004276-00-0020 |
| ON-TARGETplus Human VHL siRNA | Horizon | #L-003936-00-0020 |
| ON-TARGETplus Human IRP2 siRNA | Horizon | #L-022281-01-0020 |
| FBXL5 qPCR primers<br>Fwd - GTCTTTGGACCTCCCATGAACT<br>Rev - GGCACCAATAGACTGGCTCA | This Paper | N/A |
| HIF2α qPCR primers<br>Fwd - GCGCTAGACTCCGAGAACAT<br>Rev - TGGCCACTTACTACCTGACCCTT | This Paper | N/A |

| | | |
|---|---|---|
| VHL qPCR primers<br>Fwd - AGAGCGATGCCTCCAGGTT<br>Rev - TGACGATGTCCAGTCTCCTGTAA | This Paper | N/A |
| VEGF-A qPCR primers<br>Fwd - GGCAAAAACGAAAGCGCAAG<br>Rev - GGAGGCTCCAGGGCATTAGA | This Paper | N/A |
| GLUT1 qPCR primers<br>Fwd - GAACTCTTCAGCCAGGGTCC<br>Rev - TCACACTTGGGAATCAGCCC | This Paper | N/A |
| HIF1α qPCR primers<br>Fwd - CATAAAGTCTGCAACATGGAAGGT<br>Rev - TTTGATGGGTGAGGAATGGGTT | This Paper | N/A |
| Actin qPCR primers<br>Fwd - ACCAACTGGGACGACATGGAGAAA<br>Rev - TAGCACAGCCTGGATAGCAACGTA | This Paper | N/A |
| FBXL5 gRNA for CRISPRi:<br>GGCGCCCTTTCCTGAAGAAG | This Paper | N/A |
| Recombinant DNA | | |
| pLV hU6-sgRNA hUbC-dCas9-KRAB-T2a-Puro | Addgene | Cat. #71236 |
| psPAX2 | Addgene | Cat. #12260 |
| pMD2.G | Addgene | Cat. #12259 |
| pDONR223-VHL | Addgene | Cat. #81874 |
| Software and Algorithms | | |
| MS-STATS | https://msstats.org/ | |
| DIA-NN | @DemichevLab | |
| diann2msstats.R (MSstats_labelfree_preprocessing) | https://github.com/MiguelCos/MSstats_labelfree_preprocessing | |
| Other | | |
| EZview™ Red Anti-HA Affinity Gel | Sigma-Aldrich | Cat. #E6779 |
| EZview™ Red Anti-c-Myc Affinity Gel | Sigma-Aldrich | Cat. #E6654 |
| EZview™ Red Anti-FLAG M2 | Sigma-Aldrich | Cat. #F2426 |
| Pierce™ High Capacity Streptavidin Agarose | ThermoFisher Scientific | Cat. #20357 |
